# Five amino acid mismatches in the zinc-finger domains of CELLULOSE SYNTHASE 5 and CELLULOSE SYNTHASE 6 modulate their incorporation into cellulose synthase complexes in *Arabidopsis*

**DOI:** 10.1101/2023.10.13.562211

**Authors:** Sungjin Park, Shi-You Ding

## Abstract

*Main conclusion* Different capacities for the homodimerization of CESA5 and CESA6 are modulated by five amino acid mismatches in their zinc-finger domains and motivate discriminative incorporation of these CESAs into CSCs

Cellulose synthase 5 (CESA5) and CESA6 are known to share substantial functional overlap. In the zinc-finger domain of CESA5, there are five amino acid (AA) mismatches when compared to CESA6. These mismatches in CESA5 were replaced with their CESA6 counterparts one by one until all were replaced, generating nine engineered CESA5s. Each N-terminal enhanced yellow fluorescent protein-tagged engineered CESA5 was introduced to *prc1-1*, a *cesa6* null mutant, and resulting mutants were subjected to phenotypic analyses. We found that five single AA-replaced CESA5 proteins partially rescue the *prc1-1* mutant phenotypes to different extents. Multi-AA replaced CESA5s further rescued the mutant phenotypes in an additive manner, culminating in full recovery by CESA5^G43R+S49T+S54P+S80A+Y88F^. Investigations in cellulose content, cellulose synthase complex (CSC) motility, and cellulose microfibril organization in the same mutants support the results of the phenotypic analyses. Bimolecular fluorescence complementation assays demonstrated that the level of homodimerization in every engineered CESA5 is substantially higher than CESA5. The mean fluorescence intensity of CSCs carrying each engineered CESA5 fluctuates with the degree to which the *prc1-1* mutant phenotypes are rescued by introducing a corresponding engineered CESA5, indicating that these mismatches modulate the incorporation of both CESAs into CSCs by controlling their ability to homodimerize.

## Introduction

Cellulose is a major component of terrestrial plant cell walls, facilitating load-bearing and tensile properties as well as morphological remodeling in the cell walls (Polko and Kieber, 2019). The cellulose microfibril (CMF) comprises a bundle of *para*-crystalline (1→4)-β-D-glucan chains, making it a source of glucose molecules essential in manufacturing various industrial items, biofuels, and biomaterials (Ragauskas et al. 2006). An earlier landmark study visually confirmed, through electron microscopy, the presence of plasma membrane-bound hexameric rosette structures named cellulose synthase complexes (CSCs), from which CMFs are synthesized in vascular plants (Haigler and Brown Jr., 1986). A series of groundbreaking studies elegantly demonstrated that a CSC is comprised of three different classes of cellulose synthases (CESAs) in *Arabidopsis thaliana*. CSC assembly has been believed to occur in the Golgi body based on the presence of two Golgi-localized proteins, STELLO1 and 2, which were identified as glycosyltransferases that are associated with the CSC assembly and trafficking in the Golgi (Worden et al. 2012; Zhang et al. 2016). CSCs are transferred to the plasma membrane, where they synthesize CMFs while moving bidirectionally under the guidance of cortical microtubules (Paredez et al. 2006; Persson et al. 2007; Gutierrez et al. 2009). Recent studies demonstrated that the direction of motile CSCs is also guided by the trails of pre-existing CMFs synthesized by other CSCs in the plasma membrane (Chan and Coen, 2020; Khan and Persson, 2020).

Comprehensive genetic and biochemical studies demonstrated that CESA1 and CESA3 are indispensable for two positions in a CSC lobe and a remaining third position is occupied by one of the CESA6-like proteins to complete the CSC assembly and accomplish primary cell wall (PCW) synthesis in *Arabidopsis* (Desprez et al. 2007; Persson et al. 2007). For secondary cell wall (SCW) synthesis, CESA4, CESA7, and CESA8 are known to engage in cellulose synthesis and all these CESA isoforms have been shown to play important roles to this process (Taylor et al. 2003).

In *Arabidopsis*, CESA2, CESA5, CESA6, and CESA9 are classified as CESA6-like proteins based on their sequence similarity and functional compatibility (Desprez et al. 2007; Persson et al. 2007). In CESA6-like proteins, CESA5 and CESA6 share the closest sequence identity and phylogenetic relationship, implying their considerable functional overlap (Endler and Persson, 2011). However, exogenous expression of CESA2, CESA5, and CESA9 under the control of native *CESA6* promoter in *cesa6* null mutants resulted in only a partial rescue of their mutant phenotypes (Persson et al. 2007), indicating incomplete functional compatibility with CESA6. Previous studies unveiled that CESA2, CESA5, and CESA9 play central roles in bolstering radial cell walls during seed coat formation. CESA5 in particular is implicated in mucilage cellulose synthesis in *Arabidopsis* in concert with CESA1, CESA3, and potentially CESA10 (Sullivan et al. 2011; Griffiths et al. 2017). In addition, cellulose content in *cesa5* mutants was found to be comparable to wild type (WT) unlike substantial reduction observed in *cesa6* mutants (Bischoff et al. 2011). These findings imply that CESA2, CESA5, and CESA9 specialize in cellulose synthesis under spatiotemporally specific conditions, while CESA6 generally engages in cellulose synthesis for the PCW.

We investigated the mechanism for selecting a CESA6-like protein for the third position in CSCs with an emphasis on CESA5 and CESA6. To accomplish this goal, we generated a series of engineered CESA5 proteins by replacing amino acid (AA) mismatches in the zinc-finger domain (ZN) of CESA5 with their CESA6 counterparts one by one until all five mismatches were replaced. This was followed by multilateral analyses on the consequences of these replacements. This stepwise replacement provides a series of engineered CESA5 proteins. We found that the exogenous expression of each engineered CESA5 tagged with the N-terminal enhanced yellow fluorescent protein (EYFP) in *prc1-1*, a *cesa6* null mutant, induces a gradual rescue of its mutant phenotypes. This restoration culminates in the full recovery of WT phenotypes in *prc1-1* by introducing engineered CESA5 carrying the CESA6 counterparts in all five mismatch positions, suggesting that these mismatches in their ZNs are critical in determining their respective functional properties. Further evaluations in cellulose content, CMF organization, and CSC motility support the pattern of rescuing the mutant phenotypes in *prc1-1*. Bimolecular fluorescence complementation (BiFC) showed that the level of homodimerization in every engineered CESA5 was significantly improved when compared to WT CESA5. Furthermore, we found that the fluorescence intensity of CSCs carrying each engineered CESA5 fluctuates in accordance with degree to which the mutant phenotypes of *prc1-1* were rescued. Taken together, our findings indicate that these five mismatches in the ZNs of CESA5 and CESA6 play pivotal roles in modulating the incorporation of both CESAs into CSCs by altering their homodimerization capacity.

## Materials and methods

### Plant materials and phenotypic analysis

All *Arabidopsis* seedlings were grown on half-strength MS medium containing 1% sucrose, 0.05% 2-(N-morpholino)ethanesulfonic acid (MES), and 0.6% phytagel at pH 5.7. All light-grown seedlings used for phenotypic analyses were grown in an upright position on the MS medium plates with a photoperiod of 16 h light/8 h dark at 22°C. All dark-grown etiolated seedlings were generated under the same growth conditions in the absence of light. To perform phenotypic analyses, the total lengths of light-grown seedlings and the hypocotyl lengths of dark-grown seedlings were measured in millimeters to the second decimal place using a digital caliper. Mean and standard deviation values were calculated based on measurements from the 7-day-old light or dark-grown seedlings from each sample and used for subsequent comparative statistical analyses in a pairwise manner. For each mutant line, seeds collected from at least four independent T0 plants were used for further propagation. T3 or subsequent generations of mutant and control plant samples were employed throughout all experiments conducted in this study.

### Plasmid vectors and transgenic plants

A series of target-specific editing in the full *CESA5* CDS was carried out by overlap extension PCR. In brief, the CDS was amplified into two split fragments, whose overlapping part contains modified nucleic acid sequences that are intended to introduce target point mutations. Subsequent PCR using both split fragments as PCR templates facilitated the amplification of full CDS containing the target mutation(s). To generate the *CESA5* CDS with the multiple mutations, the same strategy as above was adopted, except using resulting full CDS amplicons from previous PCRs as a PCR template. Each target CDS was inserted in-frame downstream to the gene cassette of *CESA6* native promoter:*EYFP* in *pCAMBIA1301*. Sequences of primers used are presented in Supplementary Table S6. Sequences of all CDS inserts were confirmed by sequencing. The resultant plasmid vectors were separately transformed into GV3101 *Agrobacterium* competent cells and introduced to *prc1-1* by the floral dipping method (Clough and Bent, 2008).

### Immunoblotting

Total protein isolated from the light-grown 7-day-old seedlings of each sample using a plant protein extraction kit from Thermo Fisher Scientific (Waltham, MA, USA) was used for immunoblottings. About 50 μg of total protein from each sample was subjected to SDS-PAGE, and then proteins were transferred to a polyvinylidene fluoride (PVDF) membrane. The membrane loaded with the proteins was incubated with either anti-GFP (1:5000, Abcam) or anti-tubulin (1:5000, Abcam) primary antibody, followed by another round of incubation with horse radish peroxidase-labeled anti-rabbit IgG secondary antibody (1:10000, Abcam). Signals from target proteins were detected using an ECL substrate reagent kit.

### Cellulose content analysis

Cellulose content was measured from one-month-old plants from each sample based on a previous protocol (Kumar and Turner, 2015). 5-cm-long stems that were excised 0.5 cm above the soil level were cut in half. Two consecutive 70% ethanol incubations at 70°C were performed for 1 h each, followed by an incubation in acetone for 5 min. Stem tissues were dried overnight at 37°C and incubated in 3 mL acetic/nitric acid mixture (90% acetic acid: 70% nitric acid: water=8:1:2) and boiled for 30 min, followed by draining the mixture by aspiration. This was followed by incubation with water and acetone in order, and then dried overnight at 37°C. Each dried sample was subjected to incubation in 1 mL of 67% sulfuric acid for an hour with shaking. 20 μL of each resultant incubate was taken and diluted with 500 μL water.1 mL of 98% sulfuric acid containing 0.3% anthrone was introduced to each dilution and boiled for 5 min, followed immediately by cooling on ice. The concentration of free glucose released from crystalline CMFs from each sample was measured by colorimetric assay using a NanoDrop 2000c spectrometer (Thermo Scientific), and then cellulose content was calculated as the protocol suggests. For this analysis, stems from at least ten independently grown plants were employed for each sample.

### Atomic force microscopy

Hypocotyl tissues from the 7-day-old etiolated seedlings of each plant line were cut into pieces as small as possible in water using micro-tools under a light microscope. The resulting pieces were gently pushed to make them adhered to the surface of a poly-lysine coated micro slide. A single layer of cell wall exposing inner CMFs was selected for imaging. AFM imaging was taken using the Bruker FastScan AFM system (Billerica, MA, USA) with ScanAsyst™ imaging mode and ScanAsyst-Fluid+ probe under the same imaging settings as described previously (Park and Ding, 2020). The directionality of cellulose microfibrils in the cell walls on AFM images was analyzed using ImageJ with the directionality plugin. For each sample, at least three independently grown etiolated seedlings were used for AFM imaging.

### Laser-scanning microscopy

Time-lapse imaging by TIRF microscopy was performed using the Olympus IX73 microscope installed with cellTIRF-4Line system (Tokyo, Japan) and an ORCA-Flash4.0 C11440 digital camera from Hamamatsu Photonics (Shizuoka, Japan). All time-lapse imaging was performed as described previously (Park and Ding, 2020) at 22°C.

To take Z-stack images, confocal microscopy was performed using Nikon A1 laser-scanning confocal microscope (Tokyo, Japan) installed with photomultiplier detectors. All Z-stack images were taken as described previously (Park and Ding, 2020) with following imaging settings: −33 offset, 106 Vickers number (HV), 514 nm laser excitation at 2% laser power along with 535 nm emission collection. Z-stack images were taken every 0.5 μm from the cell surface containing the plasma membrane to subjacent ∼30 μm thick subcellular regions. 3D rendered images were generated using Nikon Element software. Both time-lapse and Z-stack images were taken on the hypocotyl epidermal cells of 4-day-old etiolated seedlings. More than five independently prepared seedlings were imaged per plant line for TIRF microscopy and three for confocal microscopy.

### CSC movement speed analysis

Using time-lapse sequences taken by TIRF microscopy, the movement speed of motile CSCs on each sample was determined by kymograph analysis using ImageJ software installed with plugins as described previously (Park and Ding, 2020). The velocity of each CSC particle was identified by calculating the travel distance of each motile CSC particle over time based on kymographs generated from time-lapse sequences. To calculate the representative mean and standard deviation of CSC movement speed for each sample, the trajectories of multiple CSC particles (*n* > 200) were analyzed. To perform kymograph analyses for each sample, more than five different time-lapse movies were taken using at least five independently prepared etiolated seedlings.

### Bimolecular fluorescence complementation (BiFC) assay

The DNA fragment covering N to VR1 of a designated *CESA* gene was individually amplified by PCR. The full CDS of each *Plasma membrane Intrinsic Protein 1* (*PIP1*) and *PIP2* in *Arabidopsis* was amplified as well. Each amplicon was initially TA-cloned into pCR8 (Thermo Fisher Scientific), and then subcloned in-frame downstream to the 35S:half-split YFP gene cassette installed in two different BiFC vectors, YFC43 and YFN43 (Addgene) using Gateway LR clonase II (Thermo Fisher Scientific). Sequences of all DNA inserts were verified by sequencing, and the sequences of primers used are presented in Supplementary Table S7. Resulting vectors were transformed separately into GV3101 Agrobacterium competent cells. To evaluate the binary interactions between the different pairs of CESA subjects, Agrobacterium cells carrying YFC43 containing the CDS insert of CESA subject 1 or YFN43 containing the CDS insert of CESA subject 2 were cultured respectively in 7 mL LB medium containing 100 mg/L kanamycin at 30°C and 220 rpm for approximately 16 h. Transformation of tobacco (*Nicotiana benthamiana*) leaves was performed according to a previous protocol (Sparkes et al., 2006). Briefly, each cell culture was centrifuged at 3000 *g* for 10 min at 22°C and resuspended in 1 mL of infiltration medium (50 mM MES, 27.7 mM D-glucose, 2 mM Na_3_PO_4_•12H_2_O, and 100 μM acetosyringone). Additional centrifugation followed at 10,000 *g* for 1 min at 22°C, and the resulting cell pellet was resuspended in 1 mL infiltration medium. The concentration of each cell resuspension was measured at OD_600_ using a NanoDrop 2000c, followed by adjusting its concentration to 2.5 at OD_600_ using infiltration medium. To produce a single inoculum for a pair of CESA subjects, 200 μL of each equilibrated resuspension was taken and mixed by vortexing. The resulting inoculum was inoculated on the underside of tobacco leaves using a syringe. For all BiFC assays, about four to five-week-old tobacco plants were employed for inoculation on their leaves. Imaging YFP signals from inoculated tobacco leaves was performed at 72 h post-inoculation using the Olympus IX73 laser-scanning microscope system installed with an ORCA-Flash 4.0 C11440 digital camera (Hamamatsu) using a 20x LUCPlanFLN (0.45 NA) objective with following settings: 514 nm laser excitation and 510-550 nm emission collection, 100 ms exposure time, and 10% laser power. For each pair of CESA subjects, more than three independently grown tobacco plants inoculated with individually prepared inoculums were used for imaging.

### Calculating the corrected total fluorescence of CSCs carrying each EYFP-CESA subject

A description of methods for time-lapse imaging by TIRF microscopy is given earlier. The first frame of each resultant time-lapse sequence was used to measure the fluorescence intensity of CSCs carrying a designated EYFP-CESA subject. For each mutant or control plant line, five independently grown etiolated seedlings were imaged and analyzed. The values of integrated density and mean background reading were measured using ImageJ. We then calculated the values of corrected total fluorescence from each image by the following formula: integrated density – (area of selected rectangular region x mean fluorescence of background readings). A pairwise comparative statistical analysis was performed based on the calculated values of corrected total fluorescence from each sample.

## Results

### The rescue of mutant phenotypes in prc1-1 by introducing engineered CESA5 proteins occurs in an additive manner

To understand their functional roles, the five AA mismatches in CESA5 were replaced with their CESA6 counterparts one by one until all the mismatches were replaced. These successive replacements generated a series of engineered CESA5 proteins, CESA5^G43R^, CESA5^S49T^, CESA5^S54P^, CESA5^S80A^, CESA5^Y88F^, CESA5^S49T+S54P^, CESA5^G43R+S49T+S54P^, CESA5^G43R+S49T+S54P+Y88F,^ and CESA5^G43R+S49T+S54P+S80A+Y88F^. All *Arabidopsis* CESAs share a similar domain structure and their ZNs are located near the N-terminal end (Fig. 1a). Fig. 1b illustrates how the AAs in the ZN of each engineered CESA5 were replaced. In the ZN of CESA5 in *Arabidopsis*, there are five AA mismatches compared to AA sequence found in the ZN of CESA6, located at Gly43, Ser49, 54, 80, and Tyr88 (Fig. 1b). Expression of each N-terminal EYFP-tagged engineered CESA5 was controlled by the *CESA6* native promoter and introduced into the *prc1-1* mutant. Phenotypic changes of *prc1-1* were evaluated in response to the exogenous expression of individual engineered CESA5 under light- or dark growth conditions. The *prc1-1* is a *cesa6* null mutant that possesses a nonsense mutation at Glu720, resulting in non-functional CESA6 and shows stunted growth under both light- and dark conditions when compared to WT (Fagard et al. 2000). EYFP-WT CESA5 and -CESA6 were also exogenously expressed separately, under the control of the same native *CESA6* promoter, in *prc1-1* and used as controls along with WT and *prc1-1*.

**Fig. 1.**
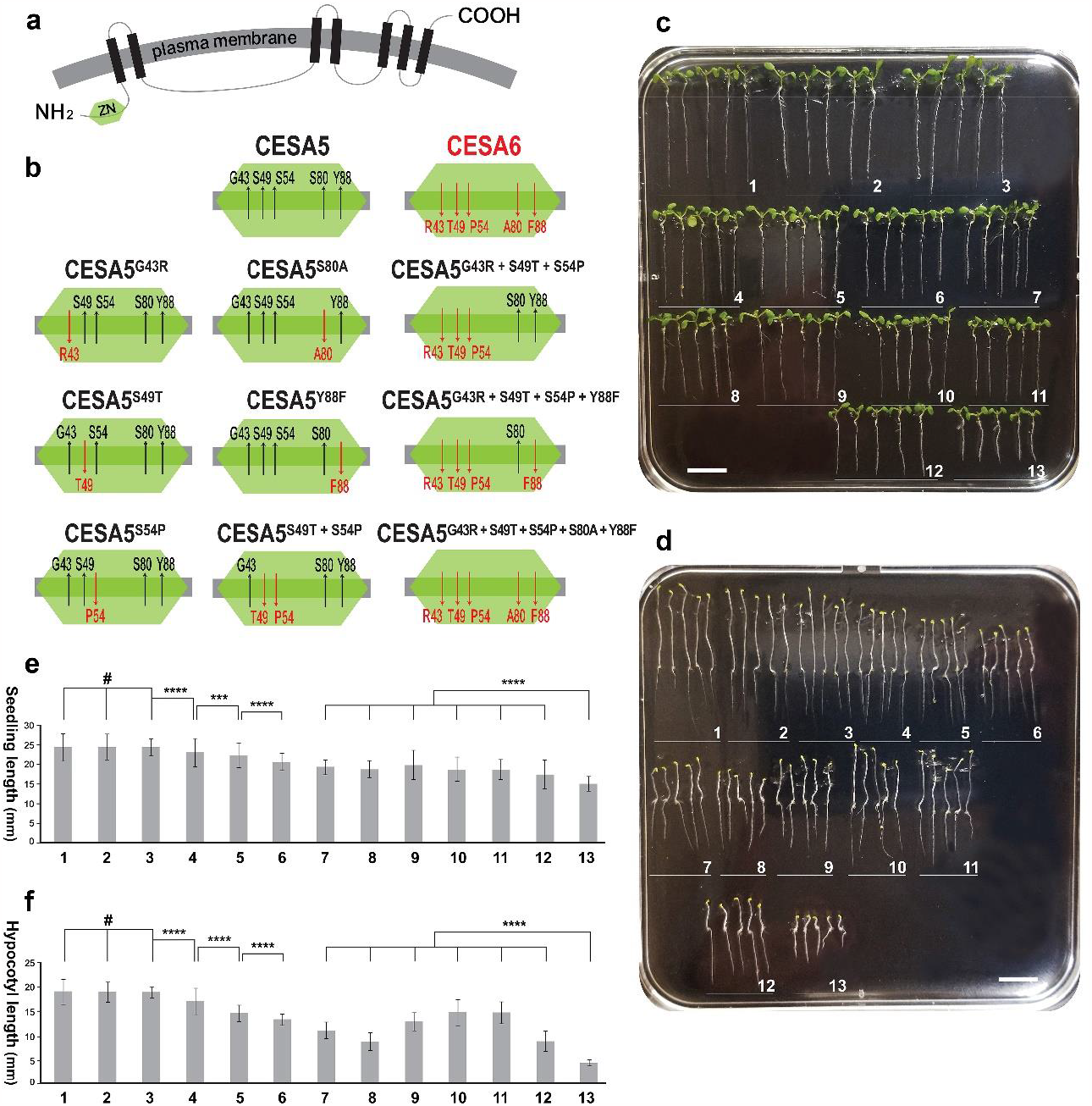
*prc1-1* mutants carrying different engineered CESA5 proteins show fluctuating phenotypic changes. (a) Schematic illustration showing a simplified structure for Arabidopsis CESAs. The green-shaded hexagon near the N-terminal end of CESA represents the ZN and black rectangular blocks indicate transmembrane domains. (b) Schematic illustrations depicting amino acid replacements in the ZN of each engineered CESA5. (c) Image showing representative 7-day-old light-grown seedlings of WT and *prc1-1* carrying each N-terminal EYFP-tagged engineered CESA5, WT CESA5, or WT CESA6, whose expression was driven by the same native *CESA6* promoter. Numbers in (c)-(f) indicate the following plant lines: 1. WT, 2. *prc1-1*+WT CESA6, 3. *prc1-1*+CESA5^G43R+S49T+S54P+S80A+Y88F^, 4. *prc1-1*+CESA5^G43R+S49T+S54P+Y88F^, 5. *prc1-1*+CESA5^G43R+S49T+S54P^, 6. *prc1-1*+CESA5^S49T+S54P^, 7. *prc1-1*+CESA5^G43R^, 8. *prc1-1*+CESA5^S49T^, 9. *prc1-1*+CESA5^S54P^, 10. *prc1-1*+CESA5^S80A^, 11. *prc1-1*+CESA5^Y88F^, 12. *prc1-1*+WT CESA5, and 13. *prc1-1*. (d) Image showing representative 7-day-old dark-grown seedlings of the same set of plant lines on c. In (c) and (d), each group of light- or dark-grown seedlings was grown together on the same half-strength MS medium plate. Scale bars = 1 cm. (e) Bar graphs represent the mean lengths of 7-day-old light-grown seedlings of different samples. Error bars stand for the standard deviations (SD) of designated samples (*n* > 364). *P*-values were calculated by two-tailed Student’s *t*-tests. (f) Bar graphs represent the mean hypocotyl lengths of 7-day-old dark-grown seedlings of different samples. Error bars stand for the SDs of designated samples (*n* > 267). Asterisks and the symbol # represent the statistical significance of analysis: \*\*\**P* < 0.0001, \*\*\*\**P* < 0.00001, and ^#^*P* > 0.4.

To probe the phenotypic consequences of introducing each engineered CESA5, WT CESA5, or CESA6 into *prc1-1*, the seedlings of all mutants and controls were grown on half-strength Murashige and Skoog (MS) medium plates for seven days under the same light-(Fig. 1c) or dark conditions (Fig. 1d), followed by phenotypic evaluation. The total lengths of 7-day-old light-grown seedlings from all mutants and controls were measured in millimeters to the second decimal place and subjected to a pairwise comparative statistical analysis (Supplementary Table S1). The mean seedling length of *prc1-1* complemented with any engineered CESA5 was found to be statistically longer than *prc1-1* or *prc1-1* complemented with WT CESA5, indicating that replacing any AA mismatches in the ZN of CESA5 with their CESA6 counterparts resulted in the functional shift of CESA5 toward CESA6 to different degrees. Based on the results of comparative statistical analysis, the mean total length of *prc1-1* seedlings complemented with CESA5^S54P^ was confirmed to be longer than any other single AA-switched CESA5. CESA5^G43R^ registered the second largest increase in mean seedling length, followed by CESA5^S49T^, CESA5^S80A^, and CESA5^Y88F^, with each exhibiting similar increases in seedling length.

The seedling lengths of *prc1-1* complemented with engineered CESA5 carrying more than one replaced AA in the mismatches were found to be further augmented in an additive manner. The mean seedling length of *prc1-1* complemented with CESA5^S49T+S54P^ is longer than *prc1-1* complemented with either CESA5^S49T^ or CESA5^S54P^. Likewise, the mean seedling length of *prc1-1* complemented with CESA5^G43R+S49T+S54P^ is longer than CESA5^S49T+S54P^, but shorter than CESA5^G43R+S49T+S54P+Y88F^. This progressive elongation of seedling lengths was found to culminate in full recovery from the stunted growth of *prc1-1* seedlings by introducing CESA5^G43R+S49T+S54P+S80A+Y88F^.The mean length of these seedlings was statistically comparable to WT or *prc1-1* complemented with CESA6, indicating the five AA mismatches collaborate on defining the functional properties of CESA5 and CESA6. The mean seedling lengths of all samples and results of the associated statistical analysis are shown in Fig. 1e.

The hypocotyl lengths of 7-day-old dark-grown *prc1-1* seedlings from the same set of plant lines were measured as well and subjected to a pairwise comparative statistical analysis (Supplementary Table S2). The progressive elongation of hypocotyls in etiolated *prc1-1* seedlings complemented with each different engineered CESA5 was observed as well. However, some minor differences were also noticed. The average hypocotyl length of etiolated seedlings of *prc1-1* complemented with CESA5^S49T^ is statistically comparable to *prc1-1* complemented with WT CESA5, suggesting that the S49T mutation does not substantially affect the hypocotyl elongation of *prc1-1* under dark conditions. Any of the other engineered CESA5s were found to prominently increase the hypocotyl elongation of etiolated *prc1-1* seedlings. Particularly, the mean hypocotyl lengths of *prc1-1* complemented with CESA5^S80A^ or CESA5^Y88F^ were found to be significantly augmented, making them statistically comparable to the mean hypocotyl length of *prc1-1* complemented with CESA5^G43R+S49T+S54P^. The mean hypocotyl lengths of all samples and results of the associated statistical analysis are given in Fig. 1f.

A previous study discovered that phytochrome B (PhyB) affects the CSC motility by regulating deterrent factors in the movement of CSCs on cortical microtubules. It has been proposed that the signaling of this regulation on CSC movement is triggered by phosphorylation on four serine residues, S122, 126, 229, and 230, in CESA5 (Bischoff et al. 2011). Another report has further confirmed that PhyB plays vital roles in sensing both temperature and light-dependent signals in plants (Legris et al. 2016). We can infer that some minor differences between light- and dark-grown seedlings may arise from this additional light-sensing regulatory system. Nevertheless, the overall pattern of rescuing the mutant phenotypes of *prc1-1* by introducing the series of engineered CESA5 proteins under light- or dark conditions is maintained in the same additive manner. Furthermore, both the stunted hypocotyl elongation and seedling growth of *prc1-1* under light- or dark conditions are fully rescued by introducing CESA5^G43R+S49T+S54P+S80A+Y88F^ when compared to *prc1-1* complemented with CESA6 or WT.

### Expression levels of all EYFP-tagged CESA proteins in prc1-1 are comparable to each other

To evaluate the expression level of N-terminal EYFP-tagged engineered CESA5, WT CESA5, or CESA6 in *prc1-1*, whose expression was governed by the same *CESA6* native promoter, two independent sessions of immunoblottings were performed using total protein from each sample. The first session of immunoblotting with anti-GFP primary antibody detected comparable amounts of target proteins, such as EYFP-tagged engineered CESA5s, WT CESA5, and CESA6, in the same *prc1-1* background at about 150 kDa, which matches the estimated size of EYFP-tagged CESA5 or CESA6. No target-sized protein band was detected in *prc1-1*, which is devoid of any EYFP-tagged CESA (Fig. 2). This result indicates that each engineered CESA5, WT CESA5, or CESA6 was exogenously expressed at a similar level in *prc1-1*. The second immunoblotting was implemented using anti-tubulin primary antibody with the same amount of total protein isolated from the same set of samples used in the first immunoblotting. This additional immunoblotting demonstrated a comparable amount of the target protein, tubulin, formed at about 50 kDa on all samples, including *prc1-1*. This confirmed that an equal amount of total protein from each sample was used for both immunoblottings.

**Fig. 2.**
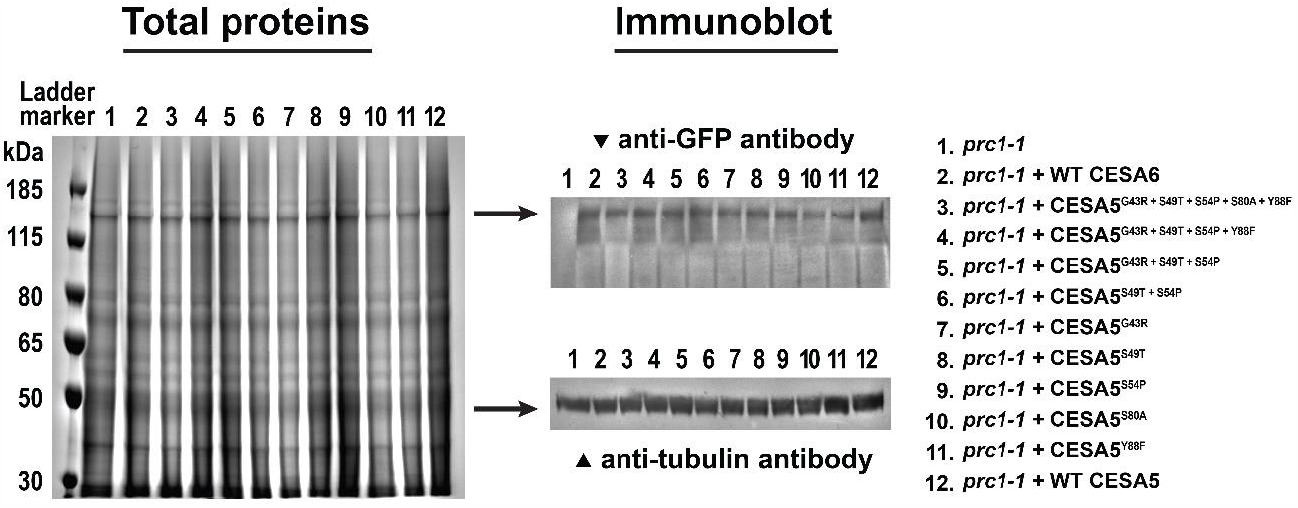
Immunoblot analyses indicate that the expression level of each EYFP-tagged engineered or WT CESA5 or CESA6 are similar to each other in *prc1-1*. Equal amounts of total protein extracted from the 7-day-old light-grown seedlings of each sample were subjected to SDS-PAGE and subsequent Coomassie blue staining (left). Two separate immunoblots were performed using anti-GFP (upper middle) or -tubulin primary antibody (bottom middle) and show that the expression of each target protein out of the same amount of total protein introduced occurred at similar levels. Identifications of samples individually numbered are specified on the right.

### The cellulose content of each mutant is in agreement with the degree of restoration of the stunted seedling growth on corresponding mutants

Cellulose content is considered a direct indication of cellulose synthesis (Kumar and Turner, 2015). In this light, the cellulose content of each sample was calculated based on the concentration of free glucose released from crystalline CMFs deposited in the cell walls of stem tissues from each sample, followed by a pairwise comparative statistical analysis. The mean cellulose contents of all samples and the results of associated pairwise statistical analysis are presented in Fig. 3 and Supplementary Table S3.

**Fig. 3.**
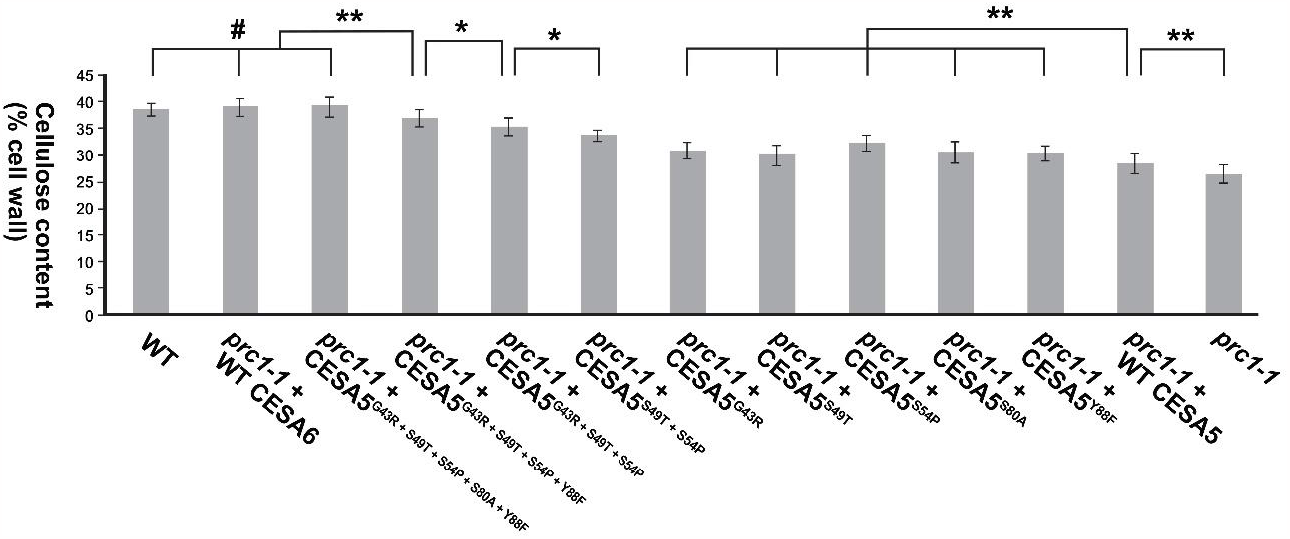
Comparative analysis of cellulose content in different mutant and control lines. Cellulose contents from different samples were calculated based on the concentration of free glucose released from crystalline CMFs in stems, followed by a comparative statistical analysis. Bar graphs and error bars respectively represent the mean cellulose contents and SDs of designated samples (*n* > 10). Fluctuation pattern in the cellulose contents of designated samples is consistent with the pattern shown in the restoration of seedling growth on the same set of samples under light conditions. *P*-values were calculated by two-tailed Student’s *t*-test. Asterisks and the symbol # denote the statistical significance of analysis: \**P* < 0.03, \*\**P* < 0.01, and ^#^*P* > 0.4.

The overall pattern of fluctuations in the cellulose contents of *prc1-1* complemented with CESA5^G43R^, CESA5^S49T^, CESA5^S54P^, CESA5^S80A^, and CESA5^Y88F^, respectively, primarily matches the pattern of rescuing stunted seedling growth in *prc1-1* seedlings complemented with a corresponding engineered CESA5 under light conditions. In addition, the cellulose contents of *prc1-1* complemented with any engineered CESA5 are statistically higher than *prc1-1* and *prc1-1* complemented with CESA5, which is a reminiscent of the case shown in the recovery of seedling growth. Differences in cellulose contents vary and correlate with the activity of cellulose synthesis conducted by CSCs carrying each different engineered CESA5. This leads to different amounts of CMF production in each sample. The cellulose content of *prc1-1* complemented with CESA5^S49T+S54P^, CESA5^G43R+S49T+S54P^, CESA5^G43R+S49T+S54P+Y88F,^ or CESA5^G43R+S49T+S54P+S80A+Y88F^ was found to increase progressively in the order listed. This is reminiscent of the pattern seen in rescuing stunted seedling growth in *prc1-1* seedlings complemented with a corresponding engineered CESA5. Likewise, the cellulose content of *prc1-1* complemented with any engineered CESA5 carrying more than one replaced mismatch with their CESA6 counterpart was verified to be higher than *prc1-1* complemented with engineered CESA5 carrying any single AA replacement as seen in rescuing the seedling growth in *prc1-1*. The escalation of cellulose content culminated in *prc1-1* complemented with CESA5^G43R+S49T+S54P+S80A+Y88F^, whose cellulose content is statistically identical to WT or *prc1-1* complemented with CESA6, suggesting that CESA5^G43R+S49T+S54P+S80A+Y88F^ is functionally equivalent to CESA6 in respect to the activity of cellulose synthesis.

### The organization of cellulose microfibrils in the cell walls of each mutant is considerably different from prc1-1

The organization of CMFs in the cell walls is fundamentally determined while they are deposited in the cell walls and by later post-modifications during cell growth (Ding and Himmel, 2006). The mode of controlling the arrangement of CMFs in the cell wall remains opaque, but it is believed that the array pattern of CSCs in the plasma membrane is directly implicated in determining the orientation of CMFs in the cell wall (Taylor et al. 2008).

We probed the near-native organization of CMFs in the cell walls of hypocotyl epidermal cells of etiolated seedlings from each sample using atomic force microscopy (AFM). A representative image of each sample is shown in Fig. 4. Based on the distributions of CMFs in different orientations (Supplementary Fig. S1), we found that the organization of CMFs in the hypocotyl epidermal cell walls of *prc1-1* complemented with any engineered CESA5 is substantially altered when compared to *prc1-1* in respect to the arrangement of CMFs. Likewise, the arrangement of CMFs in *prc1-1* carrying CESA5 or CESA6 was found to be considerably different from *prc1-1*. In *prc1-1*, the organization of CMFs appears to be relatively compact and simple because CMFs are tightly and unidirectionally deposited in the cell wall. This compact and simple arrangement of CMFs in *prc1-1* likely accounts for its overall stunted growth by constricting cell growth due to the compact and simple organization of CMFs. By contrast, the organization of CMFs in WT or *prc1-1* complemented with CESA6 appears to be more random and disordered due to multi-directed CMFs as well as larger intervals between CMFs, making it look like multilamellated mesh-like structures. Likewise, the organization of CMFs in *prc1-1* complemented with any engineered CESA5 was found to be different from *prc1-1* and appears to be similar to WT or *prc1-1* complemented with CESA6 in terms of the disordered arrangement of CMFs. Considering stunted hypocotyl elongation in etiolated *prc1-1* seedlings, the altered organization of CMFs in *prc1-1* complemented with any engineered CESA5, CESA5, or CESA6 concurs with previous reports in that the orientation of cellulose deposition is transverse to the axis of cell elongation and reoriented during cell growth accompanied by cell wall expansion over time (Lloyd and Chan, 2004). As the case shown in rescuing the mutant phenotypes and cellulose content of *prc1-1*, the organization of CMFs in *prc1-1* complemented with CESA5^G43R+S49T+S54P+S80A+Y88F^ seemingly resembles those of WT and *prc1-1* complemented with CESA6, underpinning its functional equivalency with CESA6.

**Fig. 4.**
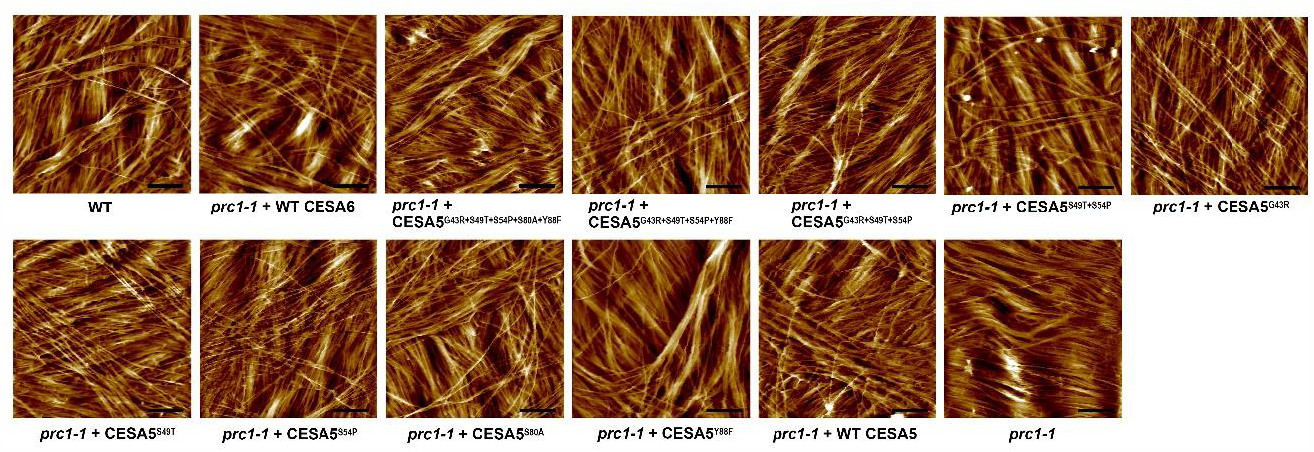
Near-native organizations of CMFs in the hypocotyl epidermal cell walls of etiolated seedlings from different plant lines show different organizational patterns. Representative AFM images taken from designated plant lines are presented. The arrangement patterns of CMFs in different samples vary considerably and show a tendency to become more complex and disordered similarly to WT by the exogenous expression of each engineered *CESA5*, WT *CESA5*, or *CESA6* in *prc1-1*. By contrast, the organization of CMFs in *prc1-1* lacks complexity and is simple and compact. Each AFM imaging was performed on an area measuring 1 μm^2^. Scale bars= 200 nm.

A previous study demonstrated that the movement and distribution patterns of motile CSCs in the plasma membrane are considerably different during PCW and SCW synthesis, and these differences determine the organization of CMFs in cell walls (Li et al. 2016). Based on given circumstances, we can infer that the engineered CESA5 proteins in CSCs, which are functionally shifted toward CESA6, make the movement and distribution patterns of active CSCs match the patterns of CSCs carrying CESA6. This in turn makes the organization of CMFs in *prc1-1* complemented with any engineered CESA5 imitate the organization of CMFs in WT or *prc1-1* complemented with CESA6.

### The velocity of motile CSCs carrying each engineered CESA5 varies in agreement with its contribution to rescuing mutant phenotypes in prc1-1

The velocity of motile CSCs in the plasma membrane is a direct indicator of the rate of cellulose synthesis. It is considered an important parameter signifying relevant changes conveyed by the mutations relating to cellulose synthesis (Woodley et al. 2018). A series of time-lapse image sequences was taken on each sample by live-cell imaging using total internal reflection fluorescence (TIRF) microscopy.

Exploiting the time-lapse image sequences, we calculated the velocities of motile CSCs carrying each engineered CESA5, WT CESA5, or CESA6 by analyzing the spatial positions of CSC particles over time and compared the motilities of CSCs on different samples. A representative kymograph for each sample is presented in Fig. 5a, and the mean velocities of CSCs carrying different EYFP-CESA subjects along with the results of associated statistical analysis are presented in Fig. 5b and Supplementary Table S4. The mean velocities of CSCs carrying CESA5^G43R^, CESA5^S49T^, CESA5^S54P^, CESA5^S80A^, or CESA5^Y88F^ were verified to be proportional to the level of rescuing hypocotyl elongation in etiolated *prc1-1* seedlings complemented with each corresponding engineered CESA5. The mean velocity of CSCs carrying WT CESA5 was found to be significantly lower than those carrying CESA6. However, it is statistically comparable with CSCs carrying CESA5^S49T^, which matches the degree of hypocotyl elongation recovery in etiolated *prc1-1* seedlings upon introducing WT CESA5 or CESA5^S49T^. Besides, the velocities of CSCs carrying CESA5^S49T+S54P^, CESA5^G43R+S49T+S54P^, CESA5^G43R+S49T+S54P+Y88F,^ or CESA5^G43R+S49T+S54P+S80A+Y88F^ were verified to increase progressively in the order listed. This concurs with the patterns shown in rescuing both mutant phenotypes and cellulose content in *prc1-1*. Furthermore, the mean velocities of CSCs carrying CESA6 or CESA5^G43R+S49T+S54P+S80A+Y88F^ were verified to be statistically identical to each other.

**Fig. 5.**
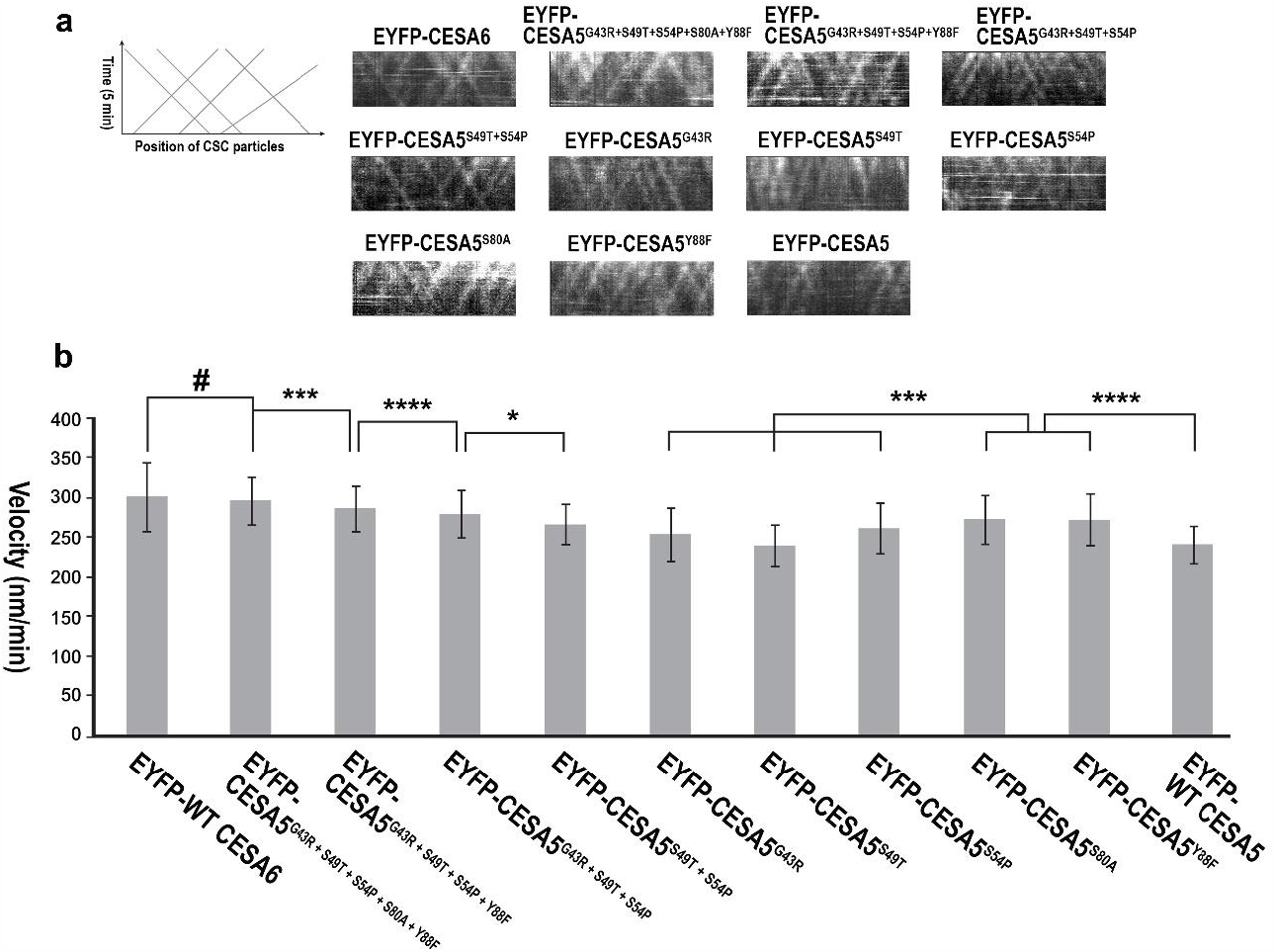
Comparative analysis of the velocity of CSCs carrying different EYFP-CESA subjects. (a) Representative kymographs of motile CSCs carrying designated EYFP-CESA subjects are presented, which were generated based on time-lapse sequences showing the motion of CSCs using ImageJ and necessary plugins. (b) Each bar graph represents the mean velocity of CSC particles carrying a designated EYFP-CESA. Each error bar represents the SD of calculated velocities of different motile CSC particles (*n* > 200) carrying a designated EYFP-CESA subject. *P*-values were calculated by two-tailed Student’s *t*-test. Asterisks and the symbol # indicate the statistical significance of analysis: \**P* < 0.011, *\*\*\*P* < 0.0003, \*\*\*\**P* < 0.00003, and ^*#*^*P* > 0.19.

Taken together, the mean velocity of motile CSCs carrying each different engineered CESA5 is elevated to different extents, reflecting a fluctuating CSC activity in *prc1-1* complemented with a corresponding engineered CESA5. Considering given circumstances, we can infer that the accelerated velocity of CSCs carrying each different engineered CESA5 is accompanied by the increased activity of cellulose synthesis in *prc1-1* complemented with a corresponding engineered CESA5. This greater cellulose synthesis likely results in an increase in cellulose content and, by extension, the rescue of mutant phenotypes in *prc1-1* accordingly.

### BiFC assays show that the homodimerization of each engineered CESA5 is substantially promoted compared to WT CESA5

BiFC assay is an advanced platform for facilitating a live-cell imaging of binary protein-protein interactions between a pair of target proteins in homologous or native species using standard molecular biology techniques and epifluorescence microscopy (Kerppola, 2008). In conventional techniques for probing protein-protein interactions, including yeast two-hybrid (Y2H), split-ubiquitin assays, and co-immunoprecipitation (co-IP), the production and post-modification of target plant proteins generally occur in heterologous systems, such as yeast and bacteria. This can negatively affect the native folding and post-modifications of target proteins, impeding original protein-protein interactions between the target proteins (Kudla and Bock, 2016). By contrast, the BiFC platform permits native or homologous systems for the target protein production and modification, which is beneficial for retaining native protein structures and properties.

We performed BiFC assays to probe the binary interaction of different CESA subjects in various pairwise combinations using the N-terminal region of each engineered CESA5, CESA1, CESA3, CESA5, or CESA6. The N-terminal region we used for BiFC assays comprises the N-terminal end (N), ZN, and variable region1 (VR1) of each CESA subject (Fig. 6a). This allows the N-terminal region of CESA5 to retain four serine residues (S122, 126, 229, and 230) in its VR1 (Fig. 6a), which are known to be crucial in the regulation of CSC motility (Bischoff et al. 2011). To minimize false results, all BiFC assays were implemented under the same stringent experimental conditions. Briefly, the same concentration of *Agrobacterium* cells carrying the expression vectors containing the coding sequence (CDS) of each CESA subject was employed for each inoculation. In this way, any false results caused by combining imbalanced or inconsistent amounts of the two CESA subjects in *Agrobacterium* cells can be reduced to monitor impartial binary interactions. In our experimental conditions, we confirmed that negative controls, a blank and a pair of plasma membrane intrinsic protein 1 (PIP1) and CESA1, do not exhibit any prominent fluorescent signals in contrast to a positive control, a pair of PIP1 and PIP2, which show substantially stronger fluorescent signals (Fig. 6b). Both positive interactions between PIP1 and PIP2 and negative interactions between PIP1 and a random CESA were demonstrated in previous studies using the same BiFC assays (Carroll et al. 2012). In addition, our BiFC results were found to coincide with the results of previous studies demonstrating binary interactions between CESA1, CESA3, CESA6, and PIPs in possible pairwise combinations (Desprez et al. 2007), indicating the reliability of our results.

**Fig. 6.**
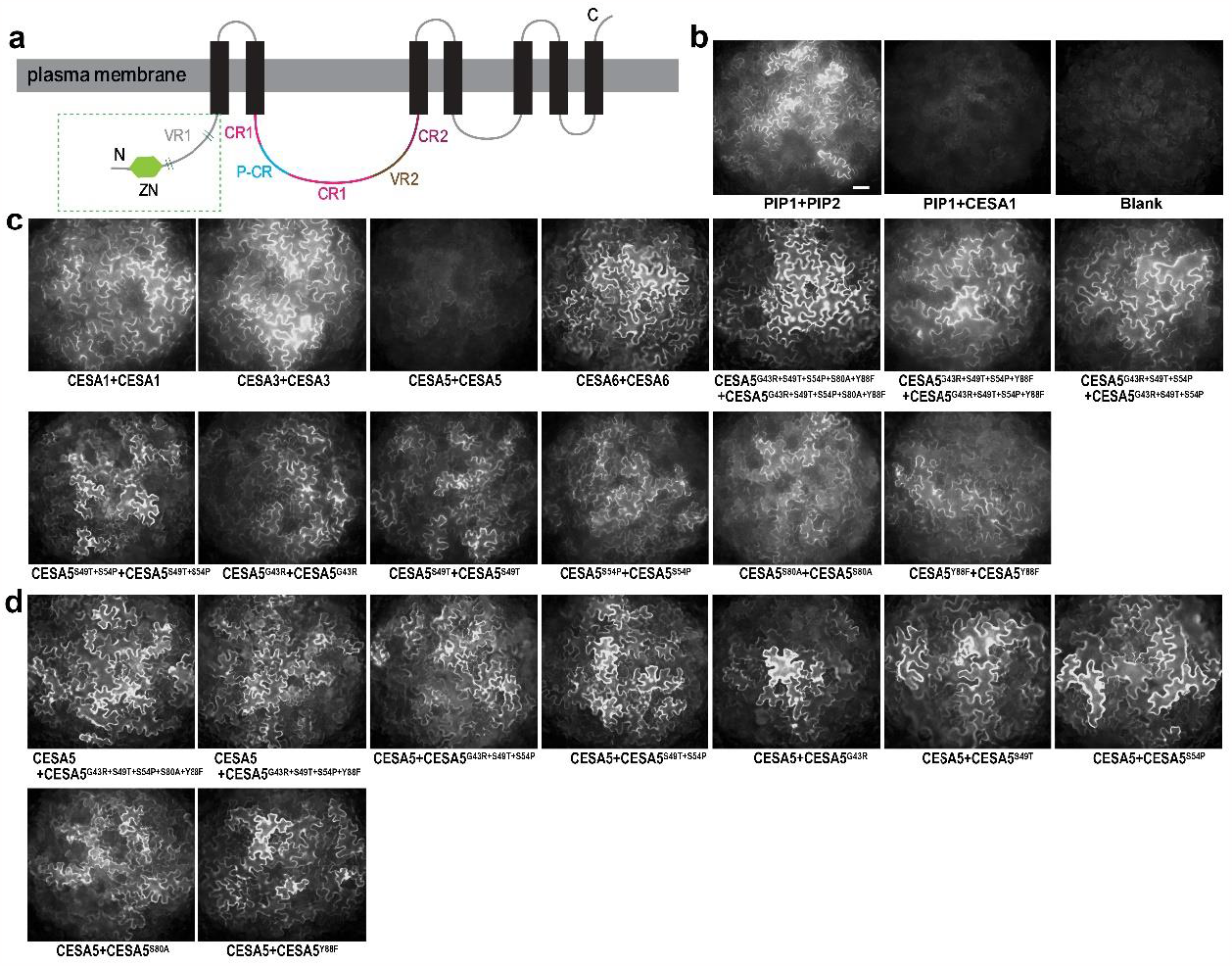
The protein-protein interactions of different pairs of primary CESA subjects. (a) Schematic diagram showing a general domain structure of *Arabidopsis* CESAs. Regions in the green dotted box covering N to VR1 were used for each CESA subject in all BiFC assays. Four short lines in the green dotted box indicate rough positions of serine residues in CESA5. Full names of abbreviations: N, N-terminal end; ZN, zinc-finger domain; TM, transmembrane domain; VR, variable region; P-CR, plant-conserved region; CR, class specific region; C, C-terminal end. (b) Representative images showing the BiFC assay results for binary protein-protein interactions in a positive control (PIP1 + PIP2) and two negative controls (PIP1 + CESA1 or blank). Scale bar = 50 μm. (c) Representative images showing homologous interactions between each primary CESA or engineered CESA5. (d) Representative images showing binary interactions between WT CESA5 and each engineered CESA5.

In all possible pairwise combinations between CESA1, CESA3, CESA5, and CESA6, we could observe significant fluorescent signals under our experimental conditions, except the homologous pairs of CESA5s (Fig. 6c). The CESA5-CESA5 pairs generate considerably weaker fluorescent signals than any other primary CESA pairs as nearly faint as the negative controls (Fig. 6, b and c). This indicates that protein-protein interaction between the same CESA5 proteins is likely weaker than any other pairs of primary CESAs, implying that attractions between the same CESA5 proteins are weaker than other primary CESA pairs. On the other hand, we could observe strong fluorescent signals in any homologous pairs of our engineered CESA5 proteins under the same conditions (Fig. 6c). This suggests that attractions between any two homologous engineered CESA5 proteins are stronger than attractions between WT CESA5 proteins, resulting in notably greater fluorescent signals between any homologous pairs of engineered CESA5s than the homologous pairs of WT CESA5. In addition, we found that any pairs of CESA5 in combination with CESA1, CESA3, or CESA6 are confirmed to produce prominent fluorescent signals (Supplementary Fig. 2b), meaning that CESA5 can intimately interact with other classes of primary CESA partners. We also confirmed that any engineered CESA5 can interact intimately with CESA6 under the same conditions (Supplementary Fig. 2a). Besides, we found that any pairs of WT CESA5 in combination with a random engineered CESA5 partner produce strong fluorescent signals as well (Fig. 6d), suggesting that our engineered CESA5 proteins behave similarly to CESA6 in respect to protein-protein interactions.

Taken together, we can infer that the five AA mismatches in the ZNs of CESA5 and CESA6 are specifically implicated in protein-protein interactions required for homodimerization. Therefore, the replacement of these mismatches in CESA5 with their CESA6 counterparts likely motivates the functional shift of CESA5 toward CESA6 by altering its capability of homodimerization.

### Mean fluorescence intensity of CSCs carrying each engineered CESA5 varies in proportion to its contribution to rescuing mutant phenotypes in prc1-1

During a series of time-lapse imaging of CSCs carrying different EYFP-CESA subjects in different samples, we found that the overall fluorescence intensity of CSCs carrying each engineered CESA5, WT CESA5, or CESA6 varies considerably under the same imaging settings and surrounding conditions. This motivated us to measure the fluorescence intensity of CSCs in different samples, followed by a comparative statistical analysis. To accomplish an accurate comparison of fluorescence between samples, every time-lapse imaging was performed on hypocotyl epidermal cells located about five to six cells below the apical hook to minimize developmental differences in target cells. In addition, to prevent any compromise in the fluorescence intensity of CSCs due to photobleaching, an optimal focal point, where CSC particles are most sharply visible, was set on a neighboring spot immediately next to a target area beforehand. This was followed by a prompt relocation to the target area while maintaining the same focal point and commencing time-lapse imaging. The measurement of fluorescence intensity of CSCs was implemented using the first frame of each time-lapse image sequence, where the fluorescence intensity of CSCs is generally the brightest. We put equal-sized rectangular boxes, whose dimensions were 10 x 20 μm, on each target area that was packed with CSC particles. The integrated density, representing the total fluorescence of CSCs in the box, was measured from five independently grown etiolated seedlings of each sample, followed by calculating a corrected total fluorescence by subtracting background fluorescence from the integrated density measured using ImageJ.

Based on the calculated values of corrected total fluorescence on different samples, a comparative statistical analysis was executed in all pairwise combinations. Representative images showing CSCs carrying each EYFP-CESA subject are presented in Fig. 7a in addition to supplementary movies showing motile CSCs carrying a corresponding EYFP-CESA (Supplementary Movies S1-S12), which all were produced under the same imaging settings and surrounding conditions. The mean fluorescence intensities of CSCs carrying different EYFP-CESA subjects, respectively, and the results of associated statistical analysis are presented in Fig. 7b and Supplementary Table S5. We verified the fluctuations of mean fluorescence intensity of CSCs carrying each EYFP-CESA subject. In addition, we found that the pattern of fluctuations in the mean fluorescence intensities of CSCs on different samples coincide with the pattern of rescuing hypocotyl elongation in etiolated *prc1-1* seedlings by introducing each corresponding EYFP-CESA. The mean fluorescence intensities of CSCs carrying EYFP-CESA5^S49T+S54P^, EYFP-CESA5^G43R+S49T+S54P^, EYFP-CESA5^G43R+S49T+S54P+Y88F^, or EYFP-CESA5^G43R+S49T+S54P+S80A+Y88F^ were found to progressively increase in the order listed, which is also consistent with the results of the other experiments conducted here. The mean fluorescence intensity of CSCs carrying EYFP-CESA5^G43R+S49T+S54P+S80A+Y88F^ is statistically comparable as EYFP-CESA6, implying that CESA5^G43R+S49T+S54P+S80A+Y88F^ can occupy the third position in a CSC as efficiently as CESA6. In this light, we can infer that this elevation of fluorescence intensity of CSCs is attributed to an increase of CSCs carrying each different engineered CESA5 compared to WT CESA5.

**Fig. 7.**
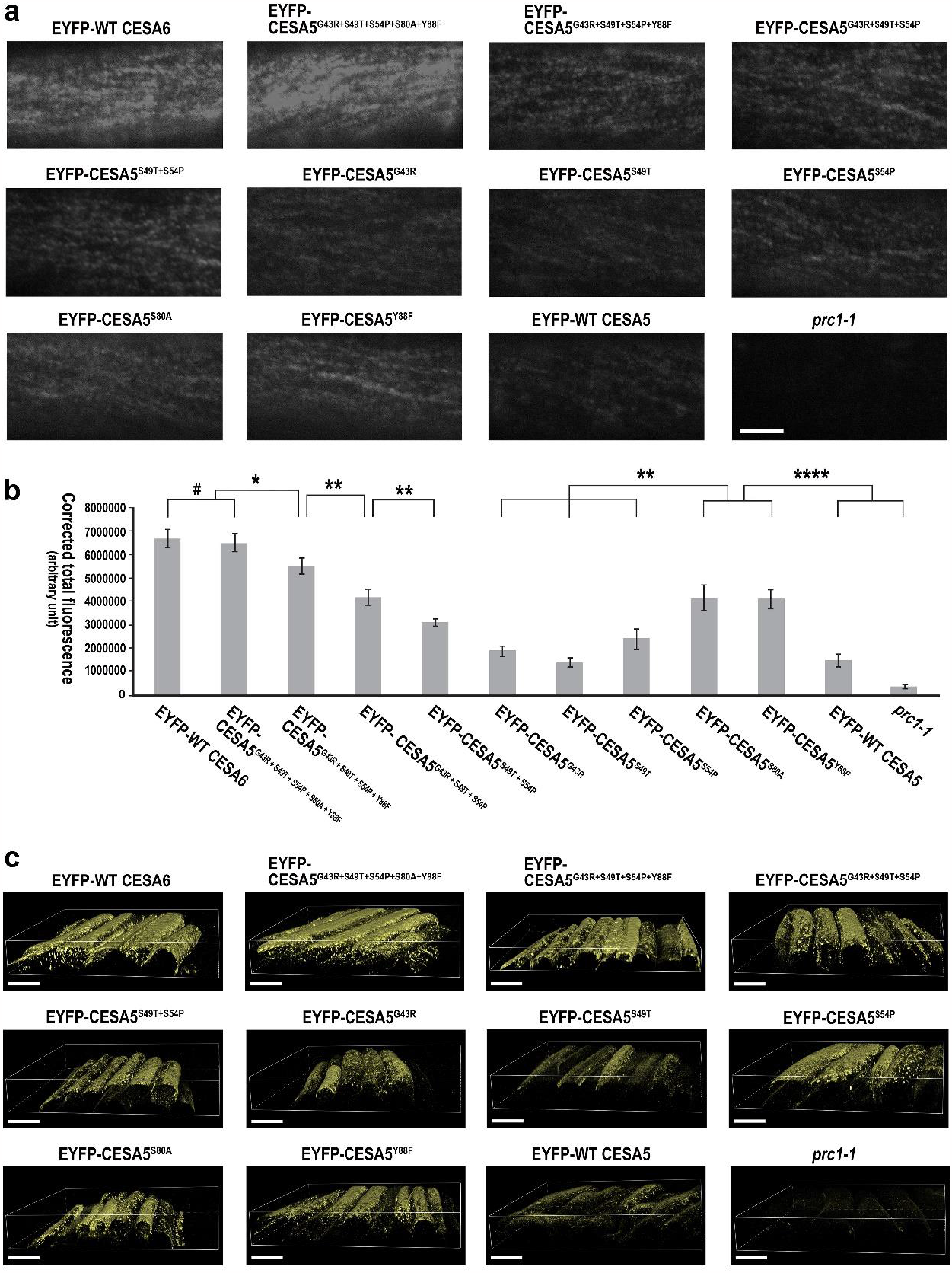
Comparative analysis of fluorescence intensity of CSCs carrying different EYFP-CESA subjects. (a) Representative images showing CSC particles carrying designated EYFP-CESA subjects. All images were taken from regions of interest with the same area (10 x 20 μm) under the same imaging settings and surrounding conditions. Scale bar = 4 μm. (b) Bar graphs and error bars respectively represent the mean values of corrected total fluorescence and SDs of designated mutant or control lines (*n* = 5). Pairwise comparative statistical analyses on the calculated values of corrected total fluorescence from different samples by two-tailed *t*-test indicate that there are significant differences between the fluorescence intensities of CSCs carrying different designated EYFP-CESA subjects. Asterisks and the symbol # indicate the statistical significance of analysis: \**P* < 0.01, \*\**P* < 0.001, \*\*\*\**P* < 0.00001, and ^#^*P* > 0.4. (c) Representative 3D rendered images showing the overall localizations and fluorescence intensities of CSCs carrying designated EYFP-CESA subjects in subcellular regions. All 3D rendered images were generated based on Z-stack images taken under the same imaging settings and surrounding conditions. Scale bars = 30 μm.

Taken together, our engineered CESA5 proteins are likely chosen preferentially for the third position in CSCs rather than WT CESA5, leading to an increase of CSCs carrying any engineered CESA5 in the plasma membrane. A direct comparison between the fluorescence intensity of EYFP-CESA5 and EYFP-CESA6 in the *prc1-1* background may not be relevant, because indigenous CESA5 proteins with no EYFP-tag exist in *prc1-1*. Hence, they compete with exogenously expressed EYFP-CESA5 for the same position in CSCs, which likely compromises the fluorescence intensity of CSCs carrying EYFP-CESA5. However, no functional native CESA6 exists in *prc1-1*, so no such internal competition occurs. On the other hand, the mean fluorescence intensity of CSCs carrying EYFP-CESA5^G43R+S49T+S54P+S80A+Y88F^ was verified to be comparable to CSCs carrying EYFP-CESA6 under the same competitive environments with endogenous CESA5 in *prc1-1*, underpinning that our EYFP-engineered CESA5 proteins are preferentially selected for the third position in a CSC rather than WT CESA5.

To further investigate the fluorescence intensity of CSCs carrying each EYFP-tagged engineered CESA5, WT CESA5, or CESA6 in subcellular regions, Z-stack images were taken from the hypocotyl epidermal cells of etiolated seedlings from each sample by confocal microscopy under the same imaging settings and surrounding conditions. Utilizing the Z-stack images, three-dimensional (3D) rendered images were computationally generated, enabling us to readily compare the subcellular distributions and overall fluorescence intensities of CSCs carrying different EYFP-CESA subjects in the same subcellular regions (Fig. 7c). Using our established imaging settings, we found that all EYFP-CESA subjects are enriched near the plasma membrane and Golgi bodies, where CSCs containing those EYFP-CESAs are localized. Furthermore, recycling of existing CSCs in the plasma membrane by clathrin-mediated endocytosis (Bashline et al. 2014) likely intensifies the enrichment of those EYFP-CESA subjects in those regions. Regarding the distribution pattern of CSCs carrying each EYFP-CESA subject, no prominent differences were found between different samples. On the other hand, we could visually observe noticeable differences in the fluorescence intensities of CSCs carrying different EYFP-CESA subjects on 3D rendered images, and the fluctuating patterns of fluorescence intensities seemingly match a pattern shown in the analytic results of fluorescence intensities of CSCs carrying corresponding CESA subjects in the plasma membrane based on the values of corrected total fluorescence.

## Discussion

Among ten CESA isomers in *Arabidopsis*, solely CESA6-like proteins have evolved to be functionally redundant and replaceable with each other, albeit partially (Persson et al. 2007). Not much is known about the underlying mechanism of selecting a CESA6-like protein for the third position in CSCs. It is unclear if the selection of a CESA6-like protein is systematically controlled or merely occurs by chance. This topic has been poorly studied despite the role it plays in the process of CSC assembly. This motivated us to conduct this research to further understand the selection mechanism of CESA6-like proteins for the third position in CSCs in *Arabidopsis*. In this study, CESA5 and CESA6 were chosen for this investigation, because they are believed to considerably share their functional properties based on their high sequence identity in spite of multiple amino acid mismatches scattered throughout different functional domains in these CESAs (Park and Ding, 2020). It is plausible that such considerable functional overlap between CESA5 and CESA6 leads to their stronger competition for the third position in CSCs when compared to other CESA6-like proteins.

Zinc fingers are protein structures known to be implicated in mediating protein-protein or protein-nucleic acid interactions (Cassandri et al. 2017). This suggests that ZNs in CESAs likely contribute to the oligomerization of CESAs in the CSC assembly. We produced a series of engineered CESA5 proteins and introduced them to *prc1-1* respectively. Consequential changes were evaluated in each mutant utilizing genetic, biochemical, and microscopic imaging techniques. In this way, we found that the functional properties of CESA5 are gradually altered and become similar to CESA6 with each successive mismatch replacement. This functional shift of CESA5 toward CESA6 was demonstrated in the progressive rescue of mutant phenotypes in *prc1-1* by introducing each engineered CESA5. This progressive rescue of mutant phenotypes in *prc1-1* culminated in full recovery by introducing CESA5^G43R+S49T+S54P+S80A+Y88F^. The fluctuations of cellulose content and CSC motility in *prc1-1* in response to the introduction of different engineered CESA5s coincide with the overall pattern of recovery in the growth of seedlings or hypocotyls of *prc1-1* under light- or dark conditions. Furthermore, the organization of CMFs deposited in the cell walls of etiolated *prc1-1* seedlings complemented with each engineered CESA5 was found to be similar to WT or *prc1-1* complemented with WT CESA6, accounting for molecular-level structural changes that are likely associated with the rescue of mutant phenotypes in *prc1-1*. In all experiments, we found that the exogenous expression of CESA5^G43R+S49T+S54P+S80A+Y88F^ in *prc1-1* results in similar impacts as introducing CESA6, implying that they are functionally equivalent. BiFC assays showed that the level of homodimerization in any engineered CESA5 was substantially improved compared to WT CESA5 and became comparable to CESA6, implying that this change in homodimerization motivates the functional shift of these engineered CESA5 proteins toward CESA6. In our experimental conditions for BiFC assays, we could not detect significant fluorescent signals between WT CESA5 proteins, but this does not mean that WT CESA5 proteins do not interact with each other and form homodimers. In our previous study, we observed stronger fluorescent signals between the same WT CESA5 proteins when an inoculum containing a higher concentration of *Agrobacterium* cells carrying the expression vectors containing the CESA5 CDS was introduced into tobacco leaves (Park and Ding, 2020).

Previous studies demonstrated the formation of CESA homodimers or homotrimers through their catalytic- and transmembrane domains under *in vitro* conditions (Purushotham et al. 2020; Qiao et al. 2021). In this study, we demonstrate that the homodimerization of CESA components, through their ZNs, is vital to facilitate the CSC assembly *in vivo* and elucidate the molecular basis behind the roles of ZNs in CESAs during CSC assembly. This suggests that the ZNs of CESAs also play fundamental roles for the CSC assembly along with catalytic- and transmembrane domains. Previous studies discovered compelling evidence demonstrating that dimerization of CESAs is crucial in establishing higher-order structures in a CSC using a combination of genetic, biochemical, and spectroscopic techniques. In cotton (*Gossypium hirsutum*), spontaneous homo- or heterodimerization of GhCESA1 and GhCESA2 through their ZNs was empirically demonstrated (Kurek et al. 2002). In addition, spontaneously formed homodimers of all SCW CESAs were found in total protein isolated from *Arabidopsis* stems by epitope tagging and successive pull-down assays (Atanassov et al. 2009). The authors propose that the homodimerization of CESA components is vital to establish further higher-order structures in CSCs. These previous findings support an idea that the greater homodimerization of each engineered CESA5 elevates the incorporation of a corresponding engineered CESA5 into CSCs compared to WT CESA5 and the resultant increase of CSCs carrying the engineered CESA5.

We found that the mean fluorescence intensity of CSCs carrying each engineered CESA5 is increased in accordance with its contribution to rescuing the hypocotyl elongation of etiolated *prc1-1* seedlings. Considering a boost in CSC motility and cellulose content in combination with the higher fluorescence intensity of CSCs by the same set of engineered CESA5s, we can infer that an increase of CSCs carrying any engineered CESA5 protein in the plasma membrane likely drives the promotion of cellulose biosynthesis. Earlier studies proposed that the propulsion of β (1→4) glucan chains synthesized by CESAs in CSCs presents a driving force for the movement of CSCs in the plasma membrane (Diotallevi and Mulder, 2007). From the information given above, we can infer that an increase of CSCs carrying any engineered CESA5 consequentially promotes the net production of glucan chains, which concomitantly accelerates the motility of CSCs. In this context, CESA5^G43R+S49T+S54P+S80A+Y88F^ likely occupies the third position in a CSC as efficiently as CESA6 due to its improved homodimerization abilities. This makes it functionally equivalent to CESA6 and thus fully compensates for the lack of a functional CESA6 in *prc1-1*.

Our study uncovers that the five AA mismatches in the ZNs of CESA5 and CESA6 are implicated in controlling their homodimerization, and consequential differences generated in their homodimerization lead to the different levels of incorporation into CSCs between CESA5 and CESA6. In short, different homodimerization between CESA5 and CESA6 drives the selective mechanism that distinguishes CESA5 from CESA6 by modulating their incorporation into CSCs and, by extension, their functional properties. Based on a high sequence similarity between CESA6-like proteins, we expect that an analogous mechanism governs the selection of CESA2 and CESA9, and further verification will follow in the future. Based on our findings, the homodimerization of CESA5 or CESA6 is vital to proceed with its oligomerization with CESA1 and CESA3 to complete the formation of a CSC. However, we do not know why homodimerized CESA5 or CESA6 facilitates their incorporation into the third position in CSCs, and how the homodimerized CESA5 or CESA6 is perceived by CESA1 and CESA3 to establish further oligomerization. Therefore, our future research will aim at uncovering the context of incorporation of homodimerized CESA6-like proteins into CSCs in concert with CESA1 and CESA3. Based on a similar approach used for evaluating CESA class specificity (Kumar et al. 2018), it is tempting to investigate the consequences of replacing the ZN of CESA1 or CESA3 with its counterpart from CESA5 or CESA6, or vice versa, with an emphasis on evaluating capacity for homodimerization, degree of incorporation into CSCs, and functional properties in resultant engineered CESAs.

Though remarkable progress has been made during the last a few decades, we are far from a comprehensive and systematic understanding in the entire process of cellulose synthesis. The advent of recent super-resolution microscopy enables us to visualize subcellular structures at sub-100-nm resolution (Schermelleh et al. 2019). Rapid technical progress raises the possibility of a direct visualization of CESA placement in a CSC in the near future. A firm grasp of the configuration of CESAs in a CSC and its assembly mechanism may enable us to manipulate cellulose synthesis and engineer the structure of CMF in the future. We hope that our findings further our understanding of the assembly mechanism for CSCs.

## Author contribution statement

SD and SP conceived the research project. SP designed and performed experiments and data analyses. SP wrote the manuscript.

## Data availability statement

The original contributions in the study are included in the article/Supplementary Material. Further inquiries can be directed to the corresponding author.

## Acknowledgements

We thank Sooyeon Park for producing all figures and illustrations, and Stephen Snyder from the School of Integrative Plant Science at Cornell University for editing and proofreading the manuscript. This work was supported by U.S. Department of Energy, Office of Science, Biological and Environmental Research Program, under Award Number DE-SC0019072.

## References

Atanassov II, Pittman JK, Turner SR (2009) Elucidating the mechanisms of assembly and subunit interaction of the cellulose synthase complex of Arabidopsis secondary cell walls. J Biol Chem 284:3833–3841.

Bashline L, Li S, Gu Y(2014) The trafficking of the cellulose synthase complex in higher plants. Ann Bot 114:1059–1067.

Bischoff V, Desprez T, Mouille G, Vernhettes S, Gonneau M, Hofte H (2011) Phytochrome regulation of cellulose synthesis in Arabidopsis. Curr Biol 21:1822–1827.

Carroll A, Mansoori N, Li S, Lei L, Vernhettes S, Viser RGF, et al. (2012) Complexes with mixed primary and secondary Cellulose synthases are functional in Arabidopsis plants. Plant Physiol 160:726–737.

Cassandri M, Smirnov A, Novelli F, Pitolli C, Agostini M, Malewicz M, et al. (2017) Zinc-finger proteins in health and disease. Cell Death Discov 3:1–12.

Chan J, Coen E (2020) Interaction between autonomous and microtubule guidance systems controls cellulose synthase trajectories. Current Biol 30:941–947.

Clough SJ, Bent AF (2008) Floral dip: a simplified method for Agrobacterium-mediated transformation of Arabidopsis thaliana. Plant J 16:735–743.

Desprez T, Juraniec M, Crowell EF, Jouy H, Pochylova Z, Parkcy F, et al. (2007) Organization of cellulose synthase complexes involved in primary cell wall synthesis in Arabidopsis thaliana. Proc Natl Acad Sci USA 104:15552–15557.

Ding S, Himmel ME (2006) The maize primary cell wall microfibril: a new model derived from direct visualization. J Agric Food Chem 54:597–606.

Diotallevi F, Mulder B (2007) The cellulose synthase complex: a polymerization driven supramolecular motor. Biophys J 92:2666–2673.

Endler A, Persson A (2011) Cellulose synthases and synthesis in Arabidopsis. Mol Plant 4:199–211.

Fagard M, Desnos T, Desprez T, Goubet F, Refregler G, Mouille G, McCann M, Rayon C, Vernhettes S, Hofte H (2000) PROCUSTE1 encodes a cellulose synthase required for normal cell elongation specifically in roots and dark-grown hypocotyls of Arabidopsis. Plant Cell 12:2409–2413.

Griffiths JS, North HM (2017) Sticking to cellulose: exploiting Arabidopsis seed coat mucilage to understand cellulose biosynthesis and cell wall polysaccharide interactions. New Phytol 214:959–966.

Gutierrez R, Lindeboom JJ, Paredez AR, Emons AMC, Ehrhardt DW (2009) Arabidopsis cortical microtubules position cellulose synthase delivery to the plasma membrane and interact with cellulose synthase trafficking compartments. Nat Cell Biol 11:797–806.

Haigler CH, Brown JRM (1986) Transport of rosettes from the Golgi apparatus to the plasma membrane in isolated mesophyll cells of Zinnia elegans during differentiation to tracheary elements in suspension culture. Protoplasma 134:111–120.

Kerppola TK (2008) Bimolecular fluorescence complementation (BiFC) analysis as a probe of protein interactions in living cells. Annu Rev Biophys 37:465–487.

Khan GA, Persson S (2020) Cell wall biology: dual control of cellulose synthase guidance. Curr Biol 30:232–234.

Kudla J, Bock R (2016) Lighting the way to protein-protein interactions: recommendations on best practices for bimolecular fluorescence complementation analyses. Plant Cell 28:1002–1008.

Kumar M, Mishra L, Carr P, Pilling M, Gardner P, Mansfield SD, Turner S (2018) Exploiting cellulose synthase (CESA) class specificity to probe cellulose microfibril biosynthesis. Plant Physiol 177:151–167.

Kumar M, Turner S (2015) Protocol: a medium-throughput method for determination of cellulose content from single stem pieces of Arabidopsis thaliana. Plant Methods 11:1–8.

Kurek I, Kawagoe Y, Jacob-Wilk D, Doblin M, Delmer D (2002) Dimerization of cotton fiber cellulose synthase catalytic subunits occurs via oxidation of the zinc-binding domains. Proc Natl Acad Sci USA 99:11109–11114.

Legris M, Klose C, Burgie ES, Rojas CC, Neme M, Hiltbrunner A, et al. (2016) Phytochrome B integrates light and temperature signals in Arabidopsis. Science 354:897–900.

Li S, Bashline L, Zheng Y, Xin X, Huang S, Kong Z, et al. (2016) Cellulose synthase complexes act in a concerted fashion to synthesize highly aggregated cellulose in secondary cell walls of plants. Proc Natl Acad Sci USA 113:11348–11353.

Lloyd C, Chan J (2004) Microtubules and the shape of plants to come. Nat Rev Mol Cell Biol 5:13–22.

Paredez AR, Somerville CR, Ehrhardt DW (2006) Visualization of cellulose synthase demonstrates functional association with microtubules. Science 312:1491–1495.

Park S, Ding S (2020) The N-terminal zinc finger of CELLULOSE SYNTHASE6 is critical in defining its functional properties by determining the level of homodimerization in Arabidopsis. Plant J 103:1826–1838.

Persson S, Paredez A, Carroll A, Palsdottir H, Doblin M, Poindexter P, et al. (2007) Genetic evidence for three unique components in primary cell-wall cellulose synthase complexes in Arabidopsis. Proc Natl Acad Sci USA 104:15566–15571.

Polko JK, Kieber JJ (2019) The regulation of cellulose biosynthesis in plants. Plant Cell 31:282–296.

Purushotham P, Ho R, Zimmer J (2020) Architecture of a catalytically active homotrimeric plant cellulose synthase complex. Science 369:1089–1094.

Qiao Z, Lampugnani ER, Yan X, Khan GA, Saw WG, Hannah P, et al. (2021) Structure of Arabidopsis CESA3 catalytic domain with its substrate UDP-glucose provides insight into the mechanism of cellulose synthesis. Proc Natl Acad Sci USA 118:2–9.

Ragauskas AJ, Williams CK, Davidson BH, Britovsek G, Cairney J, Eckert CA, et al. (2006) The path forward for biofuels and biomaterials. Science 311:484–489.

Schermelleh L, Ferrand A, Huser T, Eggeling C, Sauer M, Biehlmaier O, et al. (2019) Super-resolution microscopy demystified. Nat Cell Biol 21:72–84.

Sparkes IA, Runions J, Kearns A, Hawes C (2006) Rapid, transient expression of fluorescent fusion proteins in tobacco plants and generation of stably transformed plants. Nat Protoc 1: 2019–2025.

Sullivan S, Ralet M, Berger A, Diatloff E, Bischoff V, Gonneau M, et al. (2011) CESA5 Is required for the synthesis of cellulose with a role in structuring the adherent mucilage of Arabidopsis seeds. Plant Physiol 156:1725–1739.

Taylor NG (2008) Cellulose biosynthesis and deposition in higher plants. New Phytol 178, 239–252.

Taylor NG, Howells RM, Hutly AK, Vickers K, Turner SR (2003) Interactions among three distinct CesA proteins essential for cellulose synthesis. Proc Natl Acad Sci USA 100:1450–1455.

Woodley M, Mulvihill A, Fujita M, Wasteneys GO (2018) Exploring microtubule-dependent cellulose-synthase-complex movement with high precision particle tracking. Plants 7:1–11.

Worden N, Park E, Drakakaki G (2012) Trans-Golgi Network-An intersection of trafficking cell wall components. J Integ Plant Biol 54:875–886.

Zhang Y, Nikolovski N, Sorieul M, Vellosillo T, McFarlane HE, Dupree R, Kestern C, Schneider R, Driemeier C, Lathe R, Lampugnani E, Yu X, Ivakov A, Doblin MS, Mortimer JC, Brown SP, Persson S, Dupree P (2016) Golgi-localized STELLO proteins regulate the assembly and trafficking of cellulose synthase complexes in Arabidopsis. Nat Comm 7:1–14.

