## Supplementary Figure for "Five amino acid mismatches in the zinc-finger domains of CELLULOSE SYNTHASE 5 and CELLULOSE SYNTHASE 6 modulate their incorporation into cellulose synthase complexes in *Arabidopsis*"

**Supplementary Figures 1 and 2**

**RESEARCH ARTICLE**

### Five amino acid mismatches in the zinc-finger domains of CELLULOSE SYNTHASE 5 and CELLULOSE SYNTHASE 6 modulate their incorporation into cellulose synthase complexes in *Arabidopsis*

**Sungjin Park[1] · Shi-You Ding[1,*]**

[1]Department of Plant Biology, Michigan State University, 612 Wilson Road, East Lansing, MI 48824

### Directionality histograms

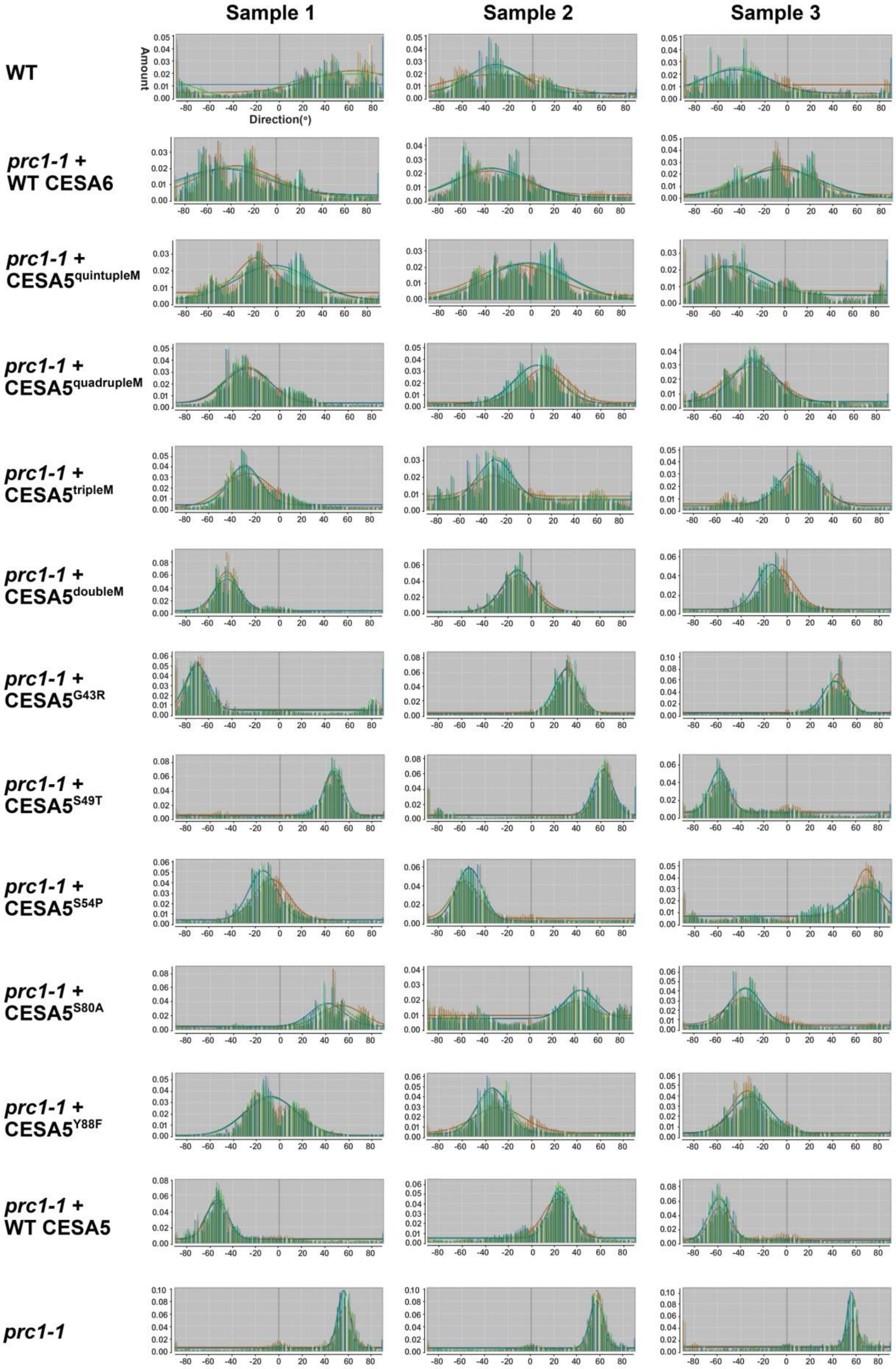

**Supplementary Fig. S1** Histograms show the distribution patterns of cellulose microfibrils laid in different orientations in the cell walls on different plant lines.

Histograms show the distribution patterns of cellulose microfibrils in the cell walls within the different ranges of relative orientation between cellulose microfibrils based on their interior angles on different plant lines. Three AFM images per plant line were analyzed using ImageJ with the directionality plugin. In the figure, quintupleM, quadrupleM, tripleM, and doubleM stand for CESA5$^{G43R+S49T+S54P+S80A+Y88F}$, CESA5$^{G43R+S49T+S54P+Y88F}$, CESA5$^{G43R+S49T+S54P}$, CESA5$^{S49T+S54P}$, respectively.

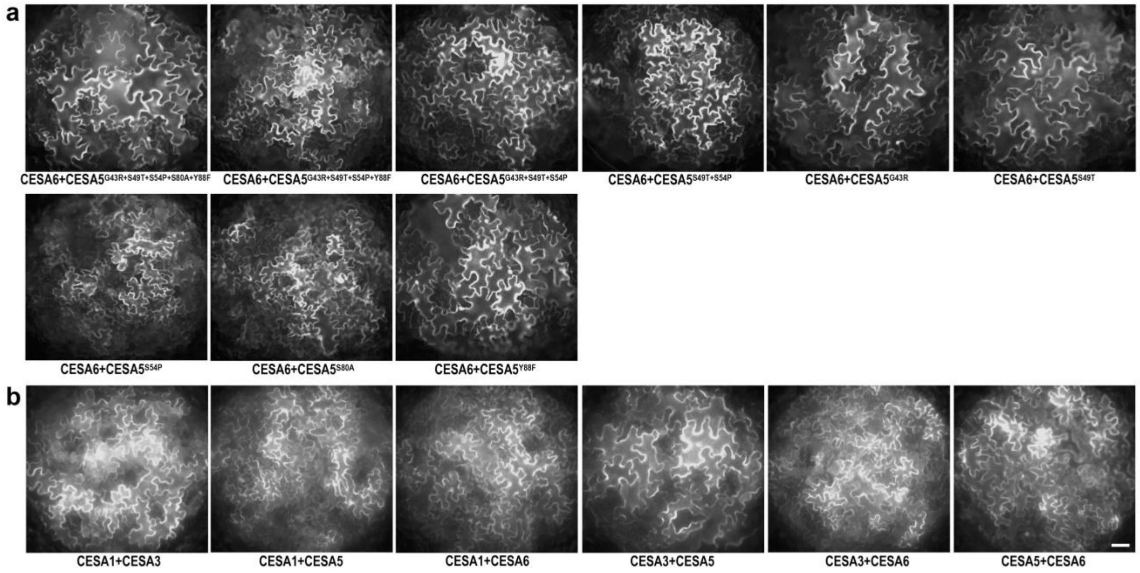


**Supplementary Fig. S2** The protein-protein interactions of different pairs of primary CESA subjects.

(a) Representative images showing binary interactions between CESA6 and each different engineered CESA5.
(b) Representative images showing heterologous interactions between primary CESAs. Scale bar = 50 µm.
