## Supplementary Table for "Five amino acid mismatches in the zinc-finger domains of CELLULOSE SYNTHASE 5 and CELLULOSE SYNTHASE 6 modulate their incorporation into cellulose synthase complexes in *Arabidopsis*"

**Supplementary Tables 1-7 and Movie Legends**

**RESEARCH ARTICLE**

### Five amino acid mismatches in the zinc-finger domains of CELLULOSE SYNTHASE 5 and CELLULOSE SYNTHASE 6 modulate their incorporation into cellulose synthase complexes in *Arabidopsis*

**Sungjin Park[1] · Shi-You Ding[1,*]**

[1]Department of Plant Biology, Michigan State University, 612 Wilson Road, East Lansing, MI 48824

**Supplementary Table S1.** The full results of pairwise comparative statistical analysis on the seedling lengths of light-grown seedlings from designated samples.

In the table, quintupleM, quadrupleM, tripleM, and doubleM stand for CESA5$^{G43R+S49T+S54P+S80A+Y88F}$, CESA5$^{G43R+S49T+S54P+Y88F}$, CESA5$^{G43R+S49T+S54P}$, CESA5$^{S49T+S54P}$, respectively. *P*-values for all pairwise combinations of samples were calculated by two-tailed *t*-test.

| *P*-value | WT | *prc1-1+* WT CESA6 | *prc1-1+* quintupleM | *prc1-1+* quadrupleM | *prc1-1+* tripleM | *prc1-1+* doubleM | *prc1-1+* CESA5$^{G43R}$ | *prc1-1+* CESA5$^{S49T}$ | *prc1-1+* CESA5$^{S54P}$ | *prc1-1+* CESA5$^{S80A}$ | *prc1-1+* CESA5$^{Y88F}$ | *prc1-1+* WT CESA5 | *prc1-1* |
|---|---|---|---|---|---|---|---|---|---|---|---|---|---|
| WT | | 0.848999 | 0.917474 | <0.00001 | <0.00001 | <0.00001 | <0.00001 | <0.00001 | <0.00001 | <0.00001 | <0.00001 | <0.00001 | <0.00001 |
| *prc1-1+* WT CESA6 | 0.848999 | | 0.712635 | <0.00001 | <0.00001 | <0.00001 | <0.00001 | <0.00001 | <0.00001 | <0.00001 | <0.00001 | <0.00001 | <0.00001 |
| *prc1-1+* quintupleM | 0.917474 | 0.712635 | | <0.00001 | <0.00001 | <0.00001 | <0.00001 | <0.00001 | <0.00001 | <0.00001 | <0.00001 | <0.00001 | <0.00001 |
| *prc1-1+* quadrupleM | <0.00001 | <0.00001 | <0.00001 | | 0.000063 | <0.00001 | <0.00001 | <0.00001 | <0.00001 | <0.00001 | <0.00001 | <0.00001 | <0.00001 |
| *prc1-1+* tripleM | <0.00001 | <0.00001 | <0.00001 | 0.000063 | | <0.00001 | <0.00001 | <0.00001 | <0.00001 | <0.00001 | <0.00001 | <0.00001 | <0.00001 |
| *prc1-1+* doubleM | <0.00001 | <0.00001 | <0.00001 | <0.00001 | <0.00001 | | <0.00001 | <0.00001 | 0.000244 | <0.00001 | <0.00001 | <0.00001 | <0.00001 |
| *prc1-1+* CESA5$^{G43R}$ | <0.00001 | <0.00001 | <0.00001 | <0.00001 | <0.00001 | <0.00001 | | 0.000332 | 0.0006 | 0.004972 | 0.001265 | <0.00001 | <0.00001 |
| *prc1-1+* CESA5$^{S49T}$ | <0.00001 | <0.00001 | <0.00001 | <0.00001 | <0.00001 | <0.00001 | 0.000332 | | <0.00001 | 0.898973 | 0.935615 | <0.00001 | <0.00001 |
| *prc1-1+* CESA5$^{S54P}$ | <0.00001 | <0.00001 | <0.00001 | <0.00001 | <0.00001 | 0.000244 | 0.0006 | <0.00001 | | <0.00001 | <0.00001 | <0.00001 | <0.00001 |
| *prc1-1+* CESA5$^{S80A}$ | <0.00001 | <0.00001 | <0.00001 | <0.00001 | <0.00001 | <0.00001 | 0.004972 | 0.898973 | <0.00001 | | 0.05152 | <0.00001 | <0.00001 |
| *prc1-1+* CESA5$^{Y88F}$ | <0.00001 | <0.00001 | <0.00001 | <0.00001 | <0.00001 | <0.00001 | 0.001265 | 0.935615 | <0.00001 | 0.05152 | | <0.00001 | <0.00001 |
| *prc1-1+* WT CESA5 | <0.00001 | <0.00001 | <0.00001 | <0.00001 | <0.00001 | <0.00001 | <0.00001 | <0.00001 | <0.00001 | <0.00001 | <0.00001 | | <0.00001 |
| *prc1-1* | <0.00001 | <0.00001 | <0.00001 | <0.00001 | <0.00001 | <0.00001 | <0.00001 | <0.00001 | <0.00001 | <0.00001 | <0.00001 | <0.00001 | |

**Supplementary Table S2.** The full results of pairwise comparative statistical analysis on the hypocotyl lengths of etiolated seedlings from designated samples.

In the table, quintupleM, quadrupleM, tripleM, and doubleM stand for CESA5$^{G43R+S49T+S54P+S80A+Y88F}$, CESA5$^{G43R+S49T+S54P+Y88F}$, CESA5$^{G43R+S49T+S54P}$, CESA5$^{S49T+S54P}$, respectively. *P*-values for all pairwise combinations of samples were calculated by two-tailed *t*-test.

| *P*-value | WT | *prc1-1*+ WT CESA6 | *prc1-1*+ quintupleM | *prc1-1*+ quadrupleM | *prc1-1*+ tripleM | *prc1-1*+ doubleM | *prc1-1*+ CESA5$^{G43R}$ | *prc1-1*+ CESA5$^{S49T}$ | *prc1-1*+ CESA5$^{S54P}$ | *prc1-1*+ CESA5$^{S80A}$ | *prc1-1*+ CESA5$^{Y88F}$ | *prc1-1*+ WT CESA5 | *prc1-1* |
|---|---|---|---|---|---|---|---|---|---|---|---|---|---|
| WT | | 0.62856 | 0.428574 | <0.00001 | <0.00001 | <0.00001 | <0.00001 | <0.00001 | <0.00001 | <0.00001 | <0.00001 | <0.00001 | <0.00001 |
| *prc1-1*+ WT CESA6 | 0.62856 | | 0.816756 | <0.00001 | <0.00001 | <0.00001 | <0.00001 | <0.00001 | <0.00001 | <0.00001 | <0.00001 | <0.00001 | <0.00001 |
| *prc1-1*+ quintupleM | 0.428574 | 0.816756 | | <0.00001 | <0.00001 | <0.00001 | <0.00001 | <0.00001 | <0.00001 | <0.00001 | <0.00001 | <0.00001 | <0.00001 |
| *prc1-1*+ quadrupleM | <0.00001 | <0.00001 | <0.00001 | | <0.00001 | <0.00001 | <0.00001 | <0.00001 | <0.00001 | <0.00001 | <0.00001 | <0.00001 | <0.00001 |
| *prc1-1*+ tripleM | <0.00001 | <0.00001 | <0.00001 | <0.00001 | | <0.00001 | <0.00001 | <0.00001 | <0.00001 | 0.312227 | 0.188145 | <0.00001 | <0.00001 |
| *prc1-1*+ doubleM | <0.00001 | <0.00001 | <0.00001 | <0.00001 | <0.00001 | | <0.00001 | <0.00001 | 0.000083 | <0.00001 | <0.00001 | <0.00001 | <0.00001 |
| *prc1-1*+ CESA5$^{G43R}$ | <0.00001 | <0.00001 | <0.00001 | <0.00001 | <0.00001 | <0.00001 | | <0.00001 | <0.00001 | <0.00001 | <0.00001 | <0.00001 | <0.00001 |
| *prc1-1*+ CESA5$^{S49T}$ | <0.00001 | <0.00001 | <0.00001 | <0.00001 | <0.00001 | <0.00001 | <0.00001 | | <0.00001 | <0.00001 | <0.00001 | 0.396657 | <0.00001 |
| *prc1-1*+ CESA5$^{S54P}$ | <0.00001 | <0.00001 | <0.00001 | <0.00001 | <0.00001 | 0.000083 | <0.00001 | <0.00001 | | <0.00001 | <0.00001 | <0.00001 | <0.00001 |
| *prc1-1*+ CESA5$^{S80A}$ | <0.00001 | <0.00001 | <0.00001 | <0.00001 | 0.312227 | <0.00001 | <0.00001 | <0.00001 | <0.00001 | | 0.913634 | <0.00001 | <0.00001 |
| *prc1-1*+ CESA5$^{Y88F}$ | <0.00001 | <0.00001 | <0.00001 | <0.00001 | 0.188145 | <0.00001 | <0.00001 | <0.00001 | <0.00001 | 0.913634 | | <0.00001 | <0.00001 |
| *prc1-1*+ WT CESA5 | <0.00001 | <0.00001 | <0.00001 | <0.00001 | <0.00001 | <0.00001 | <0.00001 | 0.396657 | <0.00001 | <0.00001 | <0.00001 | | <0.00001 |
| *prc1-1* | <0.00001 | <0.00001 | <0.00001 | <0.00001 | <0.00001 | <0.00001 | <0.00001 | <0.00001 | <0.00001 | <0.00001 | <0.00001 | <0.00001 | |

**Supplementary Table S3.** The full results of pairwise comparative statistical analysis on the cellulose contents of designated samples. In the table, quintupleM, quadrupleM, tripleM, and doubleM stand for CESA5$^{G43R+S49T+S54P+S80A+Y88F}$, CESA5$^{G43R+S49T+S54P+Y88F}$, CESA5$^{G43R+S49T+S54P}$, CESA5$^{S49T+S54P}$, respectively. *P*-values for all pairwise combinations of samples were calculated by two-tailed *t*-test.

| *P*-value | WT | *prc1-1*+ WT CESA6 | *prc1-1*+ quintupleM | *prc1-1*+ quadrupleM | *prc1-1*+ tripleM | *prc1-1*+ doubleM | *prc1-1*+ CESA5$^{G43R}$ | *prc1-1*+ CESA5$^{S49T}$ | *prc1-1*+ CESA5$^{S54P}$ | *prc1-1*+ CESA5$^{S80A}$ | *prc1-1*+ CESA5$^{Y88F}$ | *prc1-1*+ WT CESA5 | *prc1-1* |
|---|---|---|---|---|---|---|---|---|---|---|---|---|---|
| WT | | 0.609249 | 0.488313 | 0.00391 | <0.00001 | <0.00001 | <0.00001 | <0.00001 | <0.00001 | <0.00001 | <0.00001 | <0.00001 | <0.00001 |
| *prc1-1*+ WT CESA6 | 0.609249 | | 0.8517 | 0.008774 | 0.00006 | <0.00001 | <0.00001 | <0.00001 | <0.00001 | <0.00001 | <0.00001 | <0.00001 | <0.00001 |
| *prc1-1*+ quintupleM | 0.488313 | 0.8517 | | 0.008237 | 0.000046 | <0.00001 | <0.00001 | <0.00001 | <0.00001 | <0.00001 | <0.00001 | <0.00001 | <0.00001 |
| *prc1-1*+ quadrupleM | 0.00391 | 0.008774 | 0.008237 | | 0.027056 | 0.000011 | <0.00001 | <0.00001 | <0.00001 | <0.00001 | <0.00001 | <0.00001 | <0.00001 |
| *prc1-1*+ tripleM | <0.00001 | 0.00006 | 0.000046 | 0.027056 | | 0.01567 | <0.00001 | <0.00001 | 0.000264 | 0.000012 | <0.00001 | <0.00001 | <0.00001 |
| *prc1-1*+ doubleM | <0.00001 | <0.00001 | <0.00001 | 0.000011 | 0.01567 | | 0.000061 | <0.00001 | 0.02978 | 0.000284 | <0.00001 | <0.00001 | <0.00001 |
| *prc1-1*+ CESA5$^{G43R}$ | <0.00001 | <0.00001 | <0.00001 | <0.00001 | <0.00001 | 0.000061 | | 0.223919 | 0.025519 | 0.747503 | 0.469367 | 0.000865 | <0.00001 |
| *prc1-1*+ CESA5$^{S49T}$ | <0.00001 | <0.00001 | <0.00001 | <0.00001 | <0.00001 | <0.00001 | 0.223919 | | 0.010214 | 0.446524 | 0.835428 | 0.00301 | <0.00001 |
| *prc1-1*+ CESA5$^{S54P}$ | <0.00001 | <0.00001 | <0.00001 | <0.00001 | 0.000264 | 0.02978 | 0.025519 | 0.010214 | | 0.03008 | 0.033178 | 0.000019 | <0.00001 |
| *prc1-1*+ CESA5$^{S80A}$ | <0.00001 | <0.00001 | <0.00001 | <0.00001 | 0.000012 | 0.000284 | 0.747503 | 0.446524 | 0.03008 | | 0.779361 | 0.007984 | <0.00001 |
| *prc1-1*+ CESA5$^{Y88F}$ | <0.00001 | <0.00001 | <0.00001 | <0.00001 | <0.00001 | <0.00001 | 0.469367 | 0.835428 | 0.033178 | 0.779361 | | 0.006633 | <0.00001 |
| *prc1-1*+ WT CESA5 | <0.00001 | <0.00001 | <0.00001 | <0.00001 | <0.00001 | <0.00001 | 0.000865 | 0.00301 | 0.000019 | 0.007984 | 0.006633 | | 0.002058 |
| *prc1-1* | <0.00001 | <0.00001 | <0.00001 | <0.00001 | <0.00001 | <0.00001 | <0.00001 | <0.00001 | <0.00001 | <0.00001 | <0.00001 | 0.002058 | |

**Supplementary Table S4.** The full results of pairwise comparative statistical analysis on the velocities of CSCs carrying designated EYFP-CESA subjects.
In the table, quintupleM, quadrupleM, tripleM, and doubleM stand for CESA5$^{G43R+S49T+S54P+S80A+Y88F}$, CESA5$^{G43R+S49T+S54P+Y88F}$, CESA5$^{G43R+S49T+S54P}$, CESA5$^{S49T+S54P}$, respectively. *P*-values for all pairwise combinations of samples were calculated by two-tailed *t*-test.

| *P*-value | EYFP-WT CESA6 | EYFP-quintupleM | EYFP-quadrpleM | EYFP-tripleM | EYFP-doubleM | EYFP-CESA5$^{G43R}$ | EYFP-CESA5$^{S49T}$ | EYFP-CESA5$^{S54P}$ | EYFP-CESA5$^{S80A}$ | EYFP-CESA5$^{Y88F}$ | EYFP- WT CESA5 |
|---|---|---|---|---|---|---|---|---|---|---|---|
| EYFP-WT CESA6 | | 0.195119 | 0.000013 | <0.00001 | <0.00001 | <0.00001 | <0.00001 | <0.00001 | <0.00001 | <0.00001 | <0.00001 |
| EYFP-quintupleM | 0.195119 | | 0.000265 | <0.00001 | <0.00001 | <0.00001 | <0.00001 | <0.00001 | <0.00001 | <0.00001 | <0.00001 |
| EYFP-quadrpleM | 0.000013 | 0.000265 | | 0.000022 | <0.00001 | <0.00001 | <0.00001 | <0.00001 | <0.00001 | <0.00001 | <0.00001 |
| EYFP-tripleM | <0.00001 | <0.00001 | 0.000022 | | 0.010551 | <0.00001 | <0.00001 | 0.000021 | 0.719265 | 0.71597 | <0.00001 |
| EYFP-doubleM | <0.00001 | <0.00001 | <0.00001 | 0.010551 | | 0.000024 | <0.00001 | 0.048062 | 0.040454 | 0.044093 | <0.00001 |
| EYFP-CESA5$^{G43R}$ | <0.00001 | <0.00001 | <0.00001 | <0.00001 | 0.000024 | | <0.00001 | 0.020136 | <0.00001 | <0.00001 | <0.00001 |
| EYFP-CESA5$^{S49T}$ | <0.00001 | <0.00001 | <0.00001 | <0.00001 | <0.00001 | <0.00001 | | <0.00001 | <0.00001 | <0.00001 | 0.783974 |
| EYFP-CESA5$^{S54P}$ | <0.00001 | <0.00001 | <0.00001 | 0.000021 | 0.048062 | 0.020136 | <0.00001 | | 0.000179 | 0.00021 | <0.00001 |
| EYFP-CESA5$^{S80A}$ | <0.00001 | <0.00001 | <0.00001 | 0.719265 | 0.040454 | <0.00001 | <0.00001 | 0.000179 | | 0.993354 | <0.00001 |
| EYFP-CESA5$^{Y88F}$ | <0.00001 | <0.00001 | <0.00001 | 0.71597 | 0.044093 | <0.00001 | <0.00001 | 0.00021 | 0.993354 | | <0.00001 |
| EYFP- WT CESA5 | <0.00001 | <0.00001 | <0.00001 | <0.00001 | <0.00001 | <0.00001 | 0.783974 | <0.00001 | <0.00001 | <0.00001 | |

**Supplementary Table S5.** The full results of pairwise comparative statistical analysis on the values of corrected total fluorescence from CSCs carrying designated EYFP-CESA subjects.

In the table, quintupleM, quadrupleM, tripleM, and doubleM stand for CESA5$^{G43R+S49T+S54P+S80A+Y88F}$, CESA5$^{G43R+S49T+S54P+Y88F}$, CESA5$^{G43R+S49T+S54P}$, CESA5$^{S49T+S54P}$, respectively. *P*-values for all pairwise combinations of samples were calculated by two-tailed *t*-test.

| *P*-value | EYFP-WT CESA6 | EYFP-quintupleM | EYFP-quadrpleM | EYFP-tripleM | EYFP-doubleM | EYFP-CESA5$^{G43R}$ | EYFP-CESA5$^{S49T}$ | EYFP-CESA5$^{S54P}$ | EYFP-CESA5$^{S80A}$ | EYFP-CESA5$^{Y88F}$ | EYFP- WT CESA5 | *prc1-1* |
|---|---|---|---|---|---|---|---|---|---|---|---|---|
| EYFP-WT CESA6 | | 0.482655 | 0.001086 | <0.00001 | <0.00001 | <0.00001 | <0.00001 | <0.00001 | 0.000031 | <0.00001 | <0.00001 | <0.00001 |
| EYFP-quintupleM | 0.482655 | | 0.002258 | <0.00001 | <0.00001 | <0.00001 | <0.00001 | <0.00001 | 0.000044 | 0.00001 | <0.00001 | <0.00001 |
| EYFP-quadrpleM | 0.001086 | 0.002258 | | 0.000334 | <0.00001 | <0.00001 | <0.00001 | <0.00001 | 0.001545 | 0.000388 | <0.00001 | <0.00001 |
| EYFP-tripleM | <0.00001 | <0.00001 | 0.000334 | | 0.000233 | <0.00001 | <0.00001 | 0.000108 | 0.890385 | 0.749244 | <0.00001 | <0.00001 |
| EYFP-doubleM | <0.00001 | <0.00001 | <0.00001 | 0.000233 | | <0.00001 | <0.00001 | 0.00903 | 0.003504 | 0.000901 | <0.00001 | <0.00001 |
| EYFP-CESA5$^{G43R}$ | <0.00001 | <0.00001 | <0.00001 | <0.00001 | <0.00001 | | 0.005026 | 0.045022 | 0.000022 | <0.00001 | 0.037115 | <0.00001 |
| EYFP-CESA5$^{S49T}$ | <0.00001 | <0.00001 | <0.00001 | <0.00001 | <0.00001 | 0.005026 | | 0.001672 | <0.00001 | <0.00001 | 0.515814 | <0.00001 |
| EYFP-CESA5$^{S54P}$ | <0.00001 | <0.00001 | <0.00001 | 0.000108 | 0.00903 | 0.045022 | 0.001672 | | 0.000531 | 0.000225 | 0.00472 | <0.00001 |
| EYFP-CESA5$^{S80A}$ | 0.000031 | 0.000044 | 0.001545 | 0.890385 | 0.003504 | 0.000022 | <0.00001 | 0.000531 | | 0.903041 | <0.00001 | <0.00001 |
| EYFP-CESA5$^{Y88F}$ | <0.00001 | 0.00001 | 0.000388 | 0.749244 | 0.000901 | <0.00001 | <0.00001 | 0.000225 | 0.903041 | | <0.00001 | <0.00001 |
| EYFP- WT CESA5 | <0.00001 | <0.00001 | <0.00001 | <0.00001 | <0.00001 | 0.037115 | 0.515814 | 0.00472 | <0.00001 | <0.00001 | | 0.000025 |
| *prc1-1* | <0.00001 | <0.00001 | <0.00001 | <0.00001 | <0.00001 | <0.00001 | <0.00001 | <0.00001 | <0.00001 | <0.00001 | 0.000025 | |

**Supplementary Table S6.** Primers used for cloning WT or engineered *CESA5* or *CESA6* into *pCAMBIA1301*.
The nucleotide sequences of all primers used to amplify target DNA fragments for the construction of the pCAMBIA1301 plasmid vector containing the native *CESA6* promoter and the full-size coding sequence (CDS) of each engineered *CESA5*, WT *CESA5*, and *CESA6*.

| Identification | Primer DNA sequence |
|---|---|
| *CESA5* CDS 5'-end Forward primer with BglII (commonly used for all *CESA5*-based PCR products) | 5'-AATAAGATCTATGAATACTGGTGGTCGGCTCATCGCTGGTTC-3' |
| *CESA5* CDS 3'-end Forward primer with SalI (commonly used for all *CESA5*-based PCR products) | 5'-TATTGTCGACTCAAAGGCAGTCC AAGCCACATATCTCGAGGATAG-3' |
| *CESA5*$^{G43R}$ Fragment1-Reverse | 5'-CTTAGCTCAATCTCATCTCGACAGATTTGACATGTTTGTCC-3' |
| *CESA5*$^{G43R}$ Fragment2-Forward | 5'-CGAGATGAGATTGAGCTAAGTGTTGATGGAGAGTCTTTTG-3' |
| *CESA5*$^{S49T}$ Fragment1-Reverse | 5'-AAAAGACTCTCCATCAACAGTTAGCTCAATCTCATCTC-3' |
| *CESA5*$^{S49T}$ Fragment2-Forward | 5'-CTGTTGATGGAGAGTCTTTTGTGGCATGTAATGAATG-3' |
| *CESA5*$^{S54P}$ Fragment1-Reverse | 5'-CCACAAAAGGCTCTCCATCAACACTTAGCTCAATCT-3' |
| *CESA5*$^{S54P}$ Fragment2-Forward | 5'-GTGTTGATGGAGAGCCTTTTGTGGCATGTAATGAATG-3' |
| *CESA5*$^{S80A}$ Fragment1-Reverse | 5'-GCACTGAGGACAAGCTTGGTTTCCTTCTC-3' |
| *CESA5*$^{S80A}$ Fragment2-Forward | 5'-GCTTGTCCTCAGTGCAAAACTCGTTACAAGC-3' |
| *CESA5*$^{Y88F}$ Fragment1-Reverse | 5'-CCTTTGATGCGCTTGAAACGAGTTTTGCAC-3' |
| *CESA5*$^{Y88F}$ Fragment2-Forward | 5'-TCAAGCGCATCAAAGGAAGTCCAAGGG-3' |
| *CESA5*$^{S49T+S54P}$ Fragment1-Reverse | 5'-GGCTCTCCATCAACAGTTAGCTCAATCTCATCTC-3' |
| *CESA5*$^{S49T+S54P}$ Fragment2-Forward | 5'-CTGTTGATGGAGAGCCTTTTGTGGCATGTAATGAATG-3' |
| *CESA5*$^{G43R+S49T+S54P}$ Fragment1-Reverse | 5'-GTCGAGATGAGATTGAGCTAACTGTTGATGGAGAGCCTTTT-3' |
| *CESA5*$^{G43R+S49T+S54P}$ Fragment2-Forward | 5'-CTGTTGATGGAGAGCCTTTTGTGGCATGTAATGAATG-3' |
| *CESA5*$^{G43R+S49T+S54P+Y88F}$ Fragment1-Reverse | 5'-AAAAGGCTCTCCATCAACAGTTAGCTCAATCTCATCTCGAC-3' |
| *CESA5*$^{G43R+S49T+S54P+Y88F}$ Fragment2-Forward | 5'-CTGTTGATGGAGAGCCTTTTGTGGCATGTAATGAATG-3' |
| *CESA5*$^{G43R+S49T+S54P+S80A+Y88F}$ Fragment1-Reverse | 5'-GCACTGAGGACAAGCTTGGTTTCCTTCTC-3' |
| *CESA5*$^{G43R+S49T+S54P+S80A+Y88F}$ Fragment2-Forward | 5'-GCTTGTCCTCAGTGCAAAACTCGTTTCAAGC-3' |
| *CESA6* native promoter-Forward with EcoRI | 5'- GAATTCCCTATGCGTTCTGATTGCGTAATCTTAAAG-3' |
| *CESA6* native promoter-Reverse with NcoI | 5'- CCATGGATTTGTCTGAAAACAGACACAGCTAAC-3' |
| *CESA6* CDS Forward with BspMI (BglII overhang after cut) | 5'-ACCTGCTTTAGATCTATGAACACCGGTGGTCGGTTAATC-3' |
| *CESA6* CDS Forward with BspMI (SalI overhang after cut) | 5'-GTCGACAAAGCAGGTTCACAAGCAGTCTAAACCACAG-3' |

**Supplementary Table S7.** Primers used for cloning different primary *CESA* genes into BiFC vectors.

The nucleotide sequences of all primers used to amplify the CDS covering the N to VR1 of each primary CESA or engineered CESA5 as well as the full CDSs of PIP1 and PIP2 for cloning into the BiFC plasmid vectors, YFC43 or YFN43.

| Identification | Primer DNA sequence |
|---|---|
| *CESA1* N-terminal domain_Forward | 5'-ATGGAGGCCAGTGCCGGCTTG-3' |
| *CESA1* N-terminal domain_Reverse | 5'-ACGTGTATCATCAGCCATTTGGAGTTC-3' |
| *CESA3* N-terminal domain_Forward | 5'-ATGGAATCCGAAGGAGAAACCGC-3' |
| *CESA3* N-terminal domain_Reverse | 5'-CCTCGCTTCGTCATTCAGCAGAG-3' |
| WT or engineered *CESA5* N-terminal domain_Forward | 5'-ATGAATACTGGTGGTCGGCTCATC-3' |
| WT or engineered *CESA5* N-terminal domain_Reverse | 5'-CCTTCCCTCATCCATCATAGGAATATCAG-3' |
| *CESA6* N-terminal domain_Forward | 5'-ATGAACACCGGTGGTCGGTTAATC-3' |
| *CESA6* N-terminal domain_Reverese | 5'-CCTTCCCTCATCCATCATTGGAAAATCAG-3' |
| *PIP1* CDS_Forward | 5'-ATGGAAGGCAAGGAAGAAGACG-3' |
| *PIP1* CDS_Reverse | 5'-TTAGCTTCTGGACTTGAAGGGG-3' |
| *PIP2* CDS_Forward | 5'-ATGGCAAAGGATGTGGAAGC-3' |
| *PIP2* CDS-Reverse | 5'-TTAGACGTTGGCAGCACTTCTG-3' |

**Supplementary Movie S1.** Time-lapse movie showing the movement of motile CSCs carrying EYFP-WT CESA6 in an etiolated *prc1-1* seedling. The frame rate is 30 frames per second.

**Supplementary Movie S2.** Time-lapse movie showing the movement of motile CSCs carrying EYFP-CESA5$^{G43R+S49T+S54P+S80A+Y88F}$ in an etiolated *prc1-1* seedling. The frame rate is 30 frames per second.

**Supplementary Movie S3.** Time-lapse movie showing the movement of motile CSCs carrying EYFP-CESA5$^{G43R+S49T+S54P +Y88F}$ in an etiolated *prc1-1* seedling. The frame rate is 30 frames per second.

**Supplementary Movie S4.** Time-lapse movie showing the movement of motile CSCs carrying EYFP-CESA5$^{G43R+S49T+S54P}$ in an etiolated *prc1-1* seedling. The frame rate is 30 frames per second.

**Supplementary Movie S5.** Time-lapse movie showing the movement of motile CSCs carrying EYFP-CESA5$^{S49T+S54P}$ in an etiolated *prc1-1* seedling. The frame rate is 30 frames per second.

**Supplementary Movie S6.** Time-lapse movie showing the movement of motile CSCs carrying EYFP-CESA5$^{G43R}$ in an etiolated *prc1-1* seedling. The frame rate is 30 frames per second.

**Supplementary Movie S7.** Time-lapse movie showing the movement of motile CSCs carrying EYFP-CESA5$^{S49T}$ in an etiolated *prc1-1* seedling. The frame rate is 30 frames per second.

**Supplementary Movie S8.** Time-lapse movie showing the movement of motile CSCs carrying EYFP-CESA5$^{S54P}$ in an etiolated *prc1-1* seedling. The frame rate is 30 frames per second.

**Supplementary Movie S9.** Time-lapse movie showing the movement of motile CSCs carrying EYFP-CESA5$^{S80A}$ in an etiolated *prc1-1* seedling. The frame rate is 30 frames per second.

**Supplementary Movie S10.** Time-lapse movie showing the movement of motile CSCs carrying EYFP-CESA5$^{Y88F}$ in an etiolated *prc1-1* seedling. The frame rate is 30 frames per second.

**Supplementary Movie S11.** Time-lapse movie showing the movement of motile CSCs carrying EYFP-WT CESA5 in an etiolated *prc1-1* seedling. The frame rate is 30 frames per second.

**Supplementary Movie S12.** Time-lapse movie showing the movement of motile CSCs carrying no EYFP-CESA in an etiolated *prc1-1* seedling. The frame rate is 30 frames per second.
